# Ancient regulatory evolution shapes individual language abilities in present-day humans

**DOI:** 10.1101/2025.03.07.641231

**Authors:** Lucas G. Casten, Tanner Koomar, Taylor R. Thomas, Jin-Young Koh, Dabney Hofammann, Savantha Thenuwara, Allison Momany, Marlea O’Brien, Jeff C. Murray, J. Bruce Tomblin, Jacob J. Michaelson

## Abstract

Language is a defining feature of our species, yet the genomic changes enabling it remain poorly understood. Despite decades of work since *FOXP2*’s discovery, we still lack a clear picture of which regions shaped language evolution and how variation contributes to present-day phenotypic differences. Using a novel evolutionary stratified polygenic score approach in nearly 40,000 individuals, we find that Human Ancestor Quickly Evolved Regions (HAQERs) are specifically associated with language but not general cognition. HAQERs evolved before the human–Neanderthal split, giving hominins increased binding of Forkhead and Homeobox transcription factors, and show balancing selection across the past 20,000 years. Remarkably, language variants in HAQERs appear more prevalent in Neanderthals and have convergently evolved across vocal-learning mammals. Our results reveal how ancient innovations continue shaping human language.

---

Human language is one of our species’ most remarkable cognitive innovations, yet the genetic mechanisms underlying this ability remain elusive. While the human genome differs by only 1-5% from our closest primate relatives (*1–3*), these modest genetic changes enabled the evolution of our species’ unique capacity for complex language. Understanding how these relatively small genomic differences produced profound cognitive differences represents a central challenge in evolutionary genetics, with implications for language disorders, human cognitive diversity, and the origins of human-specific traits (*4*).

The discovery that mutations in *FOXP2* cause speech and language disorders provided the first clear example of a single gene with significant effects on language, reinforcing early expectations for simple genetic architectures (*5, 6*). However, *FOXP2*’s contribution to typical variation in language ability proved limited, with subsequent studies failing to find associations between common *FOXP2* variants and individual differences in language skills (*7, 8*). This limitation shifted research toward polygenic models emphasizing a large number of regulatory elements scattered throughout the genome that collectively influence language development. Genome-wide association studies have since identified numerous loci contributing to reading abilities, stuttering, rhythm, and vocabulary development, supporting a highly polygenic architecture (*9–14*). Cross-species studies have revealed that language-related traits (vocal learning and rhythm) show convergent evolution across mammalian lineages, with distributed regulatory networks rather than single genes controlling complex vocal behaviors (*15–20*). However, this polygenic model has left critical evolutionary questions unanswered: how did language-relevant regulatory elements chnage during human evolution, when did humans acquire language-promoting functions, and how do ancient evolutionary changes translate into present-day individual differences in language abilities?

To address these questions, we analyzed 65 million years of primate evolutionary history to trace the origins of language-relevant genetic variation. As part of our approach, we developed an evolutionary stratified polygenic score (ES-PGS) method that partitions genetic effects based on the evolutionary origins of their sequence context. We applied this approach across nearly 40,000 individuals with detailed language phenotyping, combined with molecular analysis, ancient DNA analysis, and cross-species genomic comparisons. This multi-modal framework enabled us to directly connect ancient genomic innovations with modern individual differences in language ability, identify neurobiological mechanisms supporting language evolution, and revealed how evolutionary trade-offs have shaped human cognitive variation.

## Results

### Dimensions of language ability

To quantify dimensions of developmental language abilities, we analyzed 17 longitudinal cognitive and language assessments administered from kindergarten through 4th grade for 350 children sampled from a community-based cohort (*21*), which we refer to as the “EpiSLI” cohort. This analysis revealed seven factors representing distinct aspects of language ability (Figure 2A). The first factor (F1), primarily driven by sentence repetition scores, represents “core language” ability. Sentence repetition strongly indicates overall language capacity, making F1 a key measure of general language competence (*22, 23*). The second factor (F2) relates to receptive vocabulary and listening comprehension, covering broad receptive language skills. The third factor (F3) specifically reflects nonverbal IQ, aligning with performance IQ at both kindergarten and 2nd grade. Factor F4 captures pre-literacy language skills, incorporating all kindergarten scores except performance IQ. Its slight correlation to F1 and F2 (*r* = 0.13 and 0.12), but not F3, suggests specificity to language (Figure 2B). Factor F5, which we call “talkativeness,” mainly reflects the number of clauses produced in a narrative task. Factor F6, based on a comprehension of concepts and directions assessment, indexes mastery of directive language (i.e., task-based instructions). Factor F7 spans a variety of assessments, with specific loading on vocabulary and grammar-related tasks, suggesting a broad, crystallized knowledge of language.

**Figure 1:**
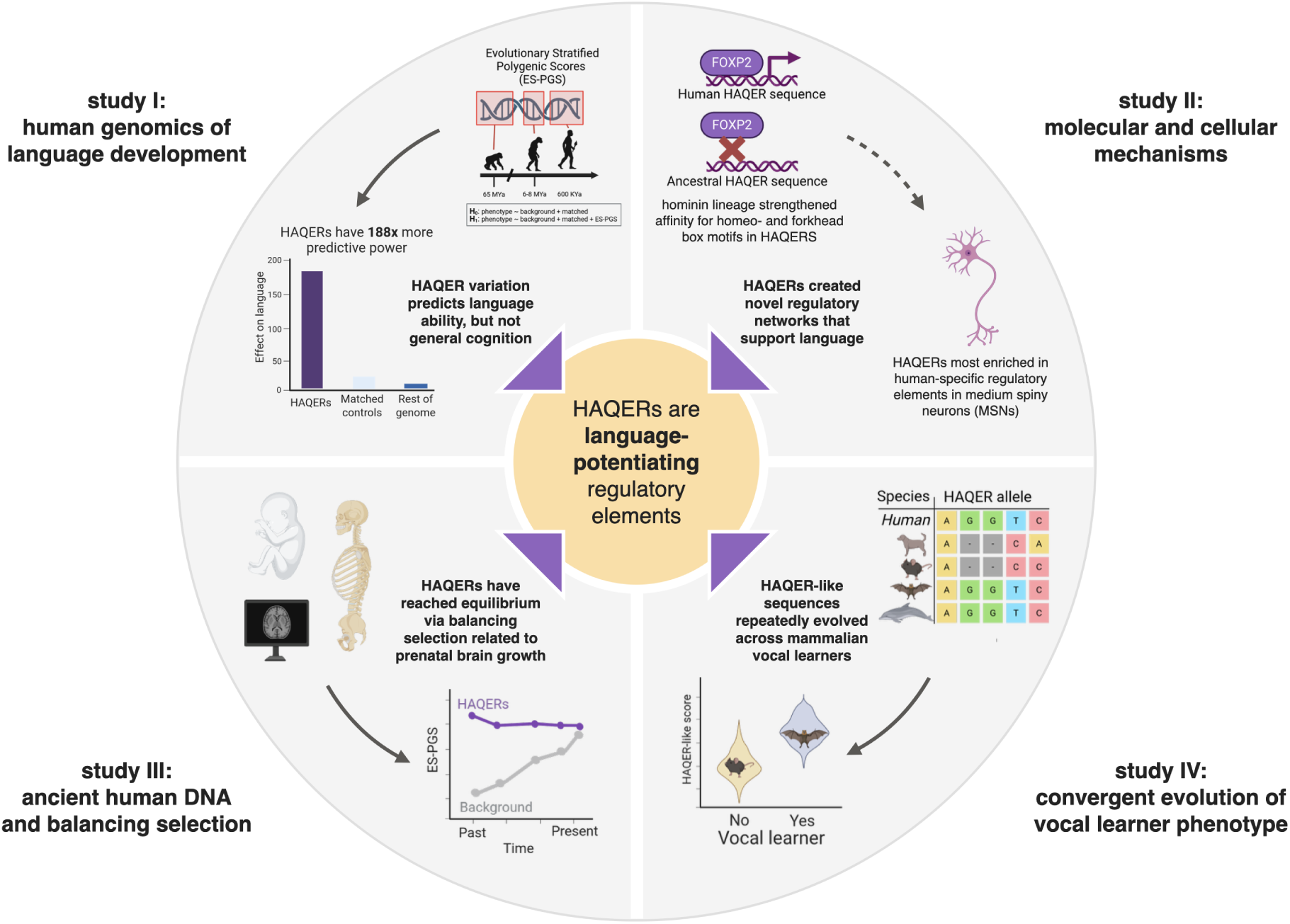
Overview of this study and key findings.

**Figure 2:**
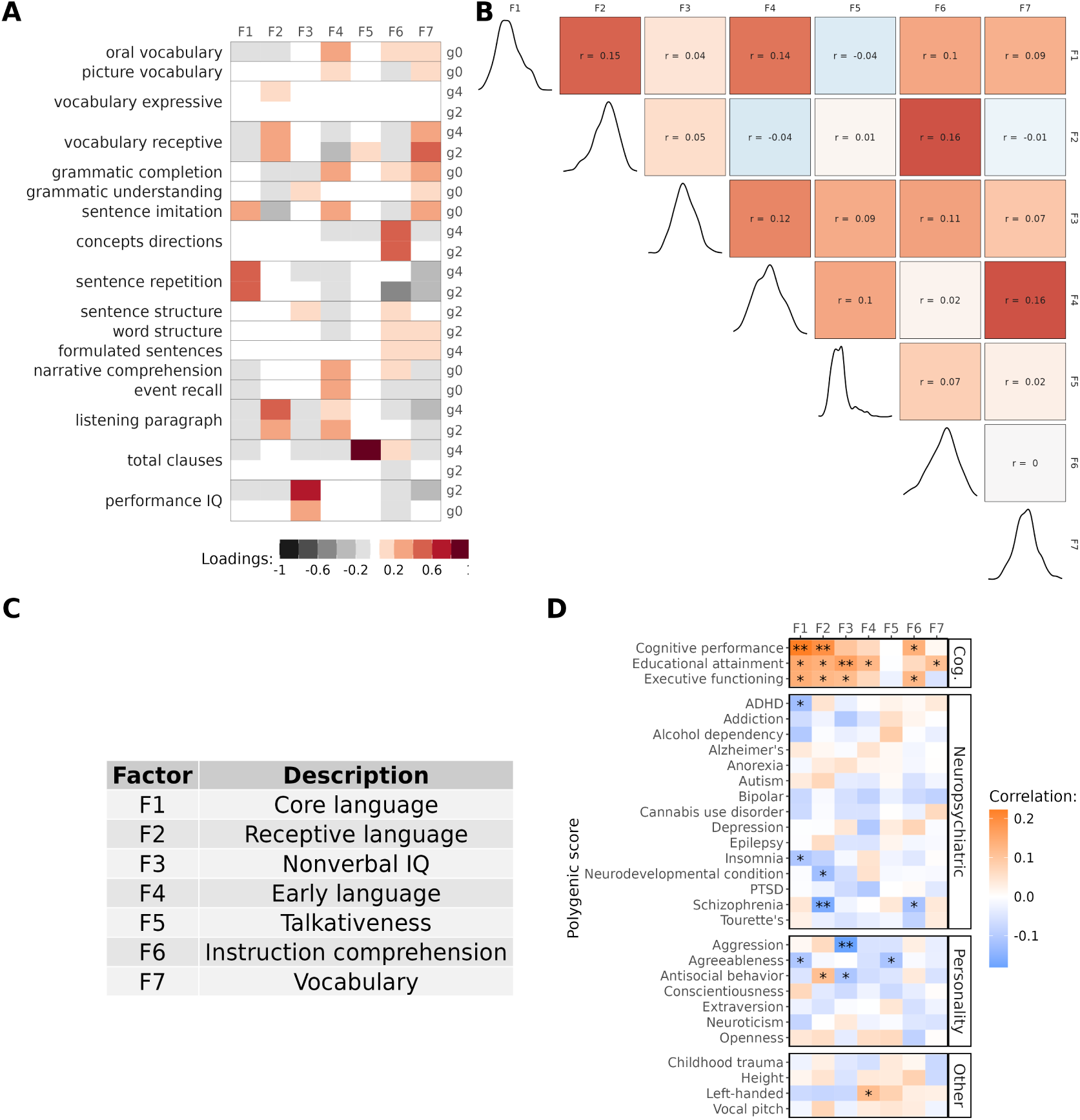
Factor loadings and genetic associations. **A** Loadings of cognitive and language assessments onto the seven language factors. g0 = Kindergarten (age 5-6), g2 = 2nd grade (age 7-8), g4 = 4th grade (age 9-10). **B** Pearson correlations for language factors (upper triangle) and distribution of each factor (diagonal). **C** Interpretations of the language factors based on their loadings. **D** Pearson correlations for each factor with genome-wide PGS. ** indicates FDR adjusted p-value *<* 0.05 and * indicates unadjusted p-value *<* 0.05.

Most of our preliminary investigation of these factors suggested that Factors 1, 2, and 3 carried the most genetic association signal (Figure 2D, Supplementary Table 2). We also find pervasive associations with F1-F3 and measures of mental health in our sample (N = 241, Figure S1, Supplementary Table 1).

### Evolutionary Stratified Polygenic Score Analysis

To investigate the genetic origins of language ability, we developed an evolutionary stratified polygenic score (ES-PGS) approach that systematically examines how genetic variants from different evolutionary periods contribute to traits. ES-PGS builds on the conceptual framework of partitioned heritability and pathway-based polygenic score methods, which have successfully partitioned genetic effects across functional genomic regions (*24–27*). However, evolutionary questions about when and how language-relevant functions emerged during human evolution require partitioning based on phylogenetic age rather than functional annotations. ES-PGS addresses this need by partitioning polygenic scores based on evolutionary origin, testing whether incorporating specific evolutionary periods significantly improves phenotypic predictions beyond what is explained by the rest of the genome and biologically matched random control regions (matched for chromosome, size, GC content, repeat content, distance to nearest gene, number of genes nearby, and overlap with promoter or coding regions). By leveraging individual-level data rather than summary statistics, ES-PGS enables direct association testing in deeply phenotyped cohorts, which is particularly valuable for specialized studies like ours with extensive language assessments.

### Human-specific genomic regions predict individual differences in language ability

We applied ES-PGS using the cognitive performance polygenic score (CP-PGS) (*28*) to trace the evolutionary origins of language abilities. This polygenic score captures broad dimensions of cognitive function, enabling valid comparisons across multiple cognitive domains (including language and nonverbal IQ). We first confirmed that the CP-PGS showed the expected associations in our EpiSLI sample, the genome-wide CP-PGS showed significant associations with both core language (F1, *r* = 0.22, FDR adjusted p-value = 0.001) and receptive language ability (F2, *r* = 0.19, FDR adjusted p-value = 0.01), (Figure 2D). Finally, we partitioned CP-PGS across five evolutionary annotations spanning approximately 65 million years of primate and human evolution, ranging from ancient primate-conserved regions to sequences differentiating modern humans from Neanderthals, to systematically trace which genomic regions and evolutionary periods contributed to different aspects of human cognition (*29–33*).

Human Ancestor Quickly Evolved Regions (HAQERs) emerged as the most compelling finding from this analysis. Despite comprising less than 0.1% of the human genome, HAQERs showed associations with four of the seven factors (F1, F2, F4, and F6, Figure 3B, Supplementary Table 3). HAQER CP-PGS demonstrated the strongest association with core language ability (*r* = 0.23, ES-PGS model *β* = 0.18, p-value = 1.2 × 10^-4^, FDR adjusted p-value = 0.004, Figures 3A-C) while showing no association with nonverbal IQ (F3, ES-PGS model *β* = 0.06, p-value = 0.19, FDR adjusted p-value = 0.61). HAQERs are largely non-coding sequences that began rapidly evolving after the human-chimpanzee split (approximately 6 million years ago) but before human-Neanderthal divergence (approximately 600,000 years ago), acquiring novel regulatory functions in the human lineage (*31, 34*).

**Figure 3:**
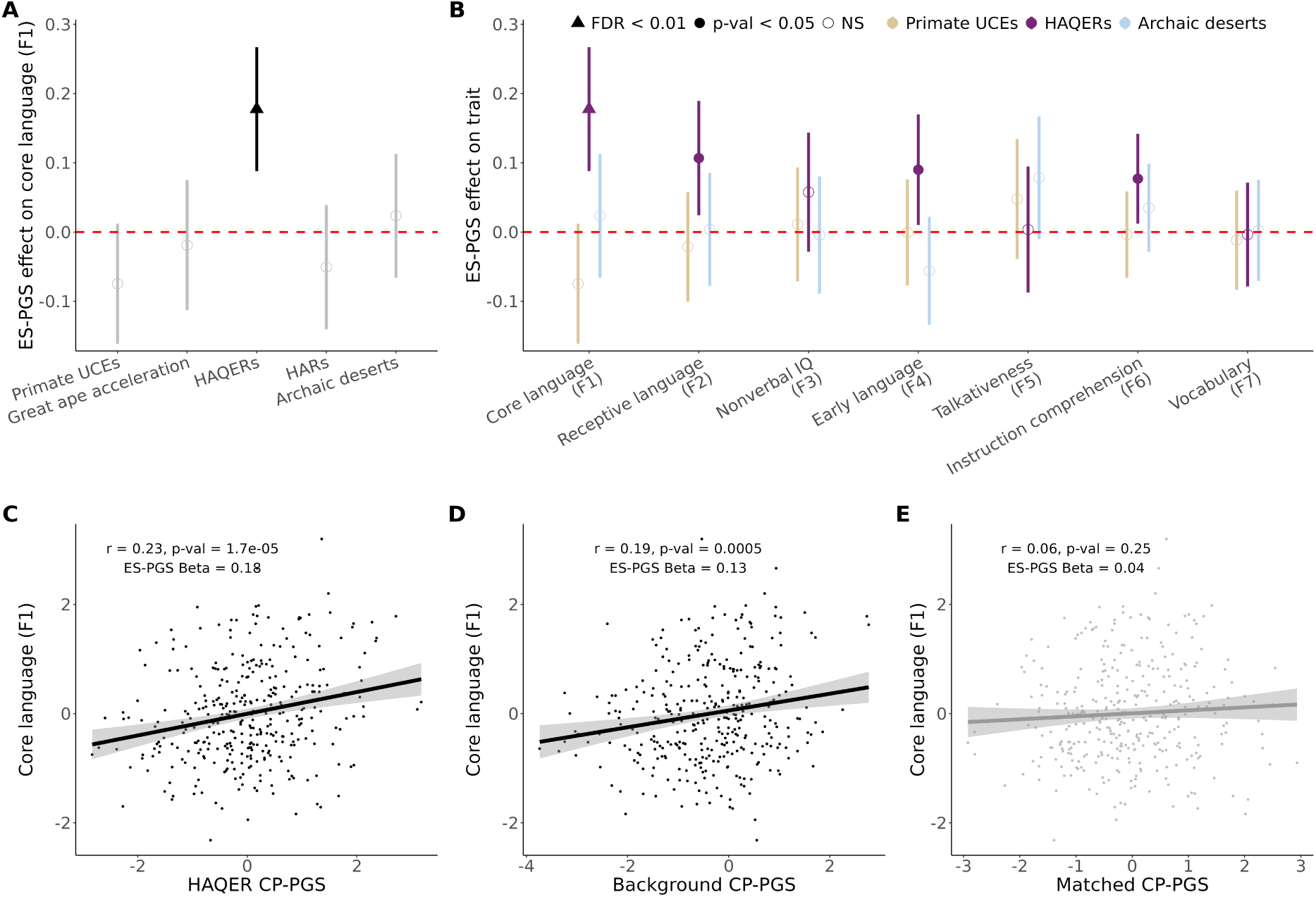
HAQERs are associated with language ability and not nonverbal IQ. **A** Comparison of evolutionary events on core language ability in EpiSLI (N = 350). Points represent the *β* provided from the ES-PGS models for each evolutionary annotation, while the ranges represent the 95% confidence interval. Solid points indicate p-value *<* 0.05. **B** Comparison of 3 evolutionary (oldest = Primate UCEs, middle = HAQERs, and youngest = archaic deserts) events on the 7 factor scores in EpiSLI (N = 350). Points represent the *β* provided from the ES-PGS models for each evolutionary annotation, while the ranges represent the 95% confidence interval. Solid points indicate p-value *<* 0.05. **C** Scatterplot of HAQER CP-PGS with core language scores (F1) in the EpiSLI sample. **D** Scatterplot of background CP-PGS with core language scores (F1) in the EpiSLI sample. **E** Scatterplot of biologically matched control regions CP-PGS with core language scores (F1) in the EpiSLI sample (matched to HAQERs).

The predictive power of HAQERs is striking. While the background and matched CP-PGS together utilized approximately 300,000 independent SNPs and explained 3.7% of variance in core language ability, adding HAQER CP-PGS (comprising only 1,763 independent SNPs) increased explained variance to 7.7%. This indicates that an average HAQER SNP carries 188 times more predictive power for language than SNPs elsewhere in the genome, with HAQERs alone explaining slightly more variance of core language scores in the EpiSLI cohort (*r*^2^ gain of 4%) than the remaining *>*99.9% of the human genome (*r*^2^ of 3.7%).

In contrast, Human Accelerated Regions (HARs), which are deeply conserved regulatory elements that acquired human-specific changes (*32, 35*), showed no comparable signal using ES-PGS (F1 *β* = -0.05, p-value = 0.27, FDR adjusted p-value = 0.7, Figure S2). This distinction suggests that human language ability emerged through novel regulatory innovations (HAQERs) rather than modifications to existing functional elements (HARs). The specific association between HAQERs and language factors but not nonverbal IQ reveals a distinct evolutionary trajectory for verbal abilities compared to general cognition. Supporting this distinction, nonverbal IQ (F3) was most strongly associated with genomic regions that underwent rapid changes across all great apes (*30*) (ES-PGS *β* = 0.13, p-value = 0.004, FDR adjusted p-value = 0.07, Figure S2).

### HAQERs influence language ability across multiple cohorts and variant types

We validated HAQER effects on language across multiple independent cohorts using both common and rare genetic variants. In the SPARK autism dataset (N *>* 30,000) (*36*), HAQER CP-PGS predicted verbal language capability (”Able to talk using short phrases or sentences”, ES-PGS *β* = 0.05, p-value = 0.008, N = 29,266) and language disorder diagnoses in parents without autism (model improvement p-value = 7.9 × 10^−5^, N = 713) but not psychiatric conditions (model improvement p-value = 0.58, N = 713), confirming language specific effects (Figure 4A-B, Supplementary Table 5). Clinical records showed HAQER CP-PGS is associated with verbal IQ (ES-PGS *β* = 2.08, p-value = 0.022, N = 620) but not nonverbal IQ (ES-PGS *β* = 0.59, p-value = 0.49, N = 620, Figure S3), further supporting specificity.

**Figure 4:**
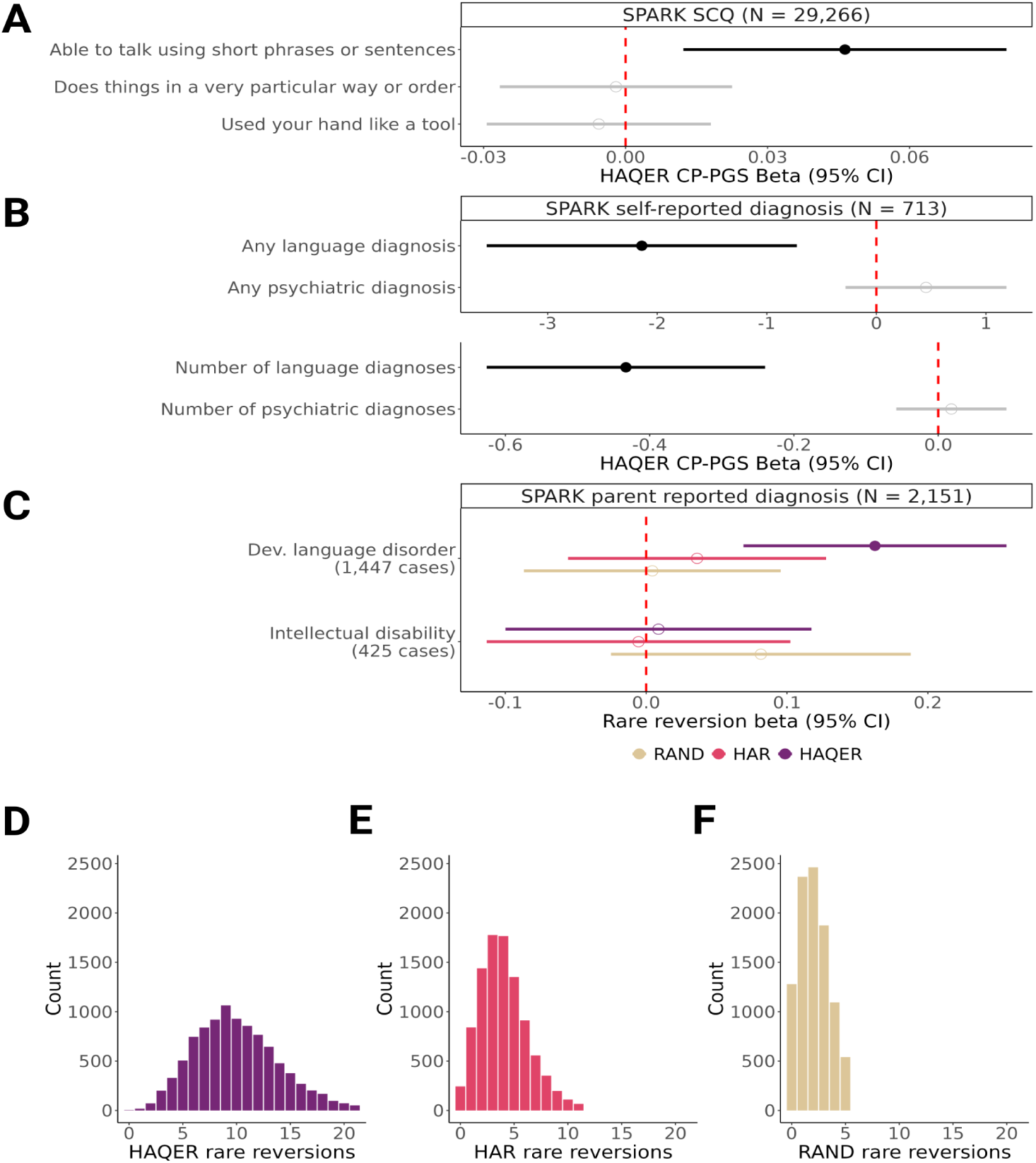
Large-scale validation of HAQERs association with language ability. **A** Points represent the *β* provided from the ES-PGS models for the HAQER CP-PGS on Social Communication Questionnaire (SCQ) items in SPARK, while the ranges represent the 95% confidence interval. Solid points indicate p-value *<* 0.05. **B** Points represent the *β* provided from the ES-PGS models for the HAQER CP-PGS on self-reported language and psychiatric diagnosis in SPARK, while the ranges represent the 95% confidence interval. Solid points indicate p-value *<* 0.05. **C** Points represent the *β* provided from the regression models for the rare reversions within 10Kb of HAQERs, HARs, or RAND (random matched) sequences, while the ranges represent the 95% confidence interval. Solid points indicate p-value *<* 0.05. **D-F** Distributions of rare reversions counts from the SPARK whole genome sequencing data within 10Kb of the following regions: HAQERs (D), HARs (E), and random sequence (RAND, F).

To test HAQER effects through an orthogonal approach independent of our ES-PGS method, we analyzed rare genetic variation in SPARK whole genome sequencing data (N *>* 2,000). We examined rare “reversions”, variants that revert from the human-specific version to their human-chimp ancestral state, reasoning that if HAQERs evolved to support human language, reversions should impair language function, which would support the observed polygenic score associations. Individuals carrying more reversions in HAQERs showed increased likelihood of developmental language disorder (*β* = 0.16, p-value = 6.5 × 10^−4^) and delayed language developmental milestones, but no association with age started walking or intellectual disability (Figure 4C, Supplementary Table 6). Notably, HAQERs showed higher rates of reversions compared to HARs and matched random sequences (Figure 4D-F).

Further independent validation in the ABCD developmental cohort (*37*) showed the HAQER CP-PGS is associated with the Rey Auditory Verbal Learning Test performance, a measure of spoken word recall (ES-PGS *β* = 0.24, p-value = 0.048, N = 5,625), but not reading or vocabulary tasks from the NIH Toolbox (Supplementary Table 7). This suggests HAQERs have preferential effects on vocal communication over written language. These converging results across cohorts and variant types strongly support that genetic variation within HAQERs have a significant and specific effect on spoken language abilities in contemporary humans.

### HAQERs evolved stronger binding affinity for language-relevant transcription factors

To investigate the molecular mechanisms underlying HAQERs’ association with language development, we analyzed how rare genetic variants affect transcription factor binding sites in these regions. We compared two classes of rare variants in the EpiSLI cohort: hominin-chimpanzee ancestral allele reversions versus other rare variants, using position weight matrices to quantify how these variants alter predicted transcription factor binding affinity. By comparing the effects of reversions with other rare variants, we could detect systematic evolutionary changes in hominin-specific transcription factor binding associated with language phenotypes.

In HAQERs, hominin-gained transcription factor motif binding showed significant correlation with individual core language ability (N = 350 individuals, *β* = 0.14, p-value = 5.6×10^−4^, Figure 5A), indicating that hominins evolved increased transcription factor binding in these regions and this binding enhances language performance. In contrast, regions under sequence conservation with human-specific changes (HARs, *β* = 0.01, p-value = 1, Figure 5B) or neutral evolution (RAND sequences, *β* = 0, p-value = 1, Figure 5C) showed no relationship between motif integrity and language ability, highlighting HAQERs’ unique and systematic selection for regulatory function during hominin evolution.

**Figure 5:**
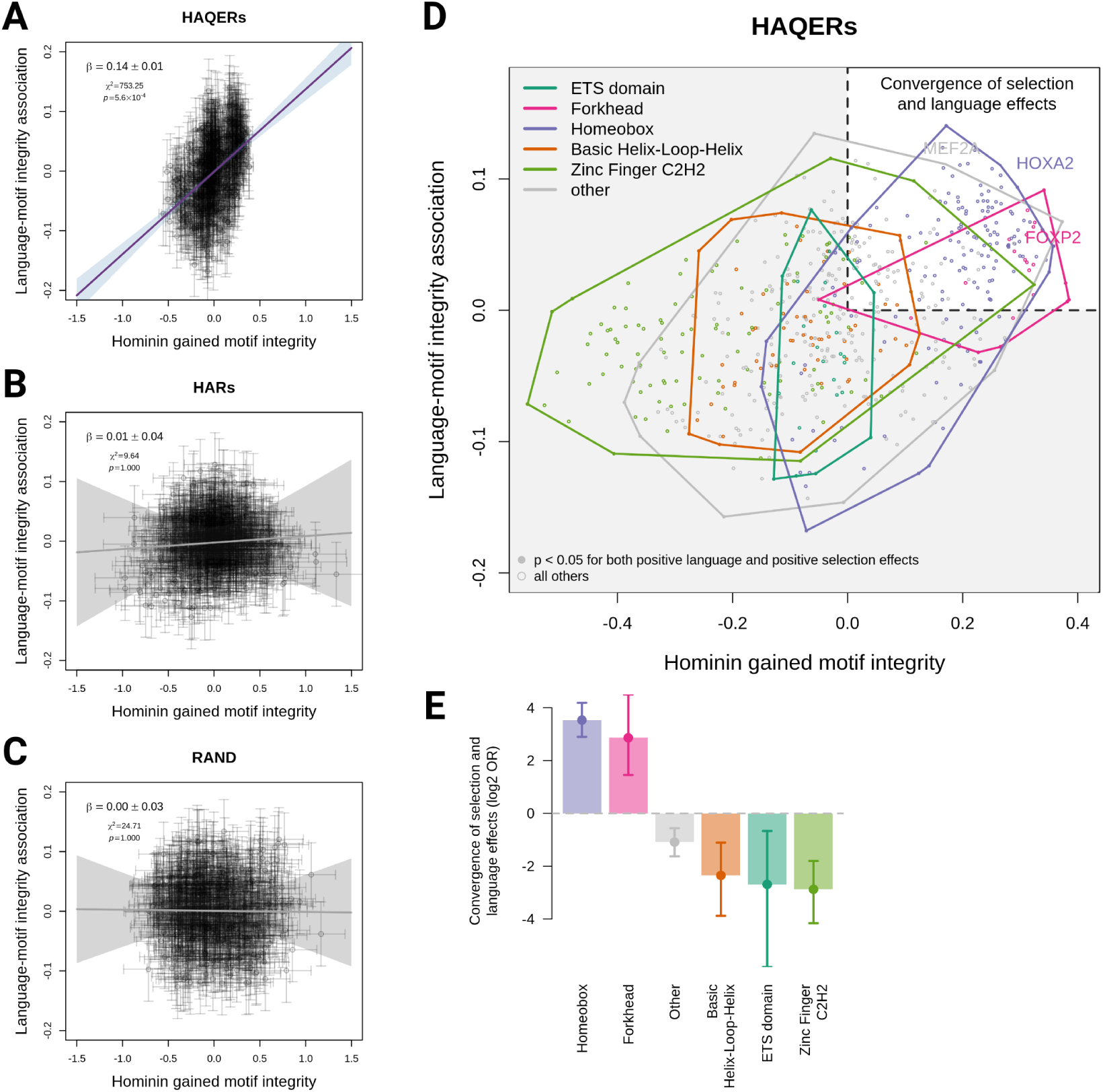
Hominin gained transcription factor binding in HAQERs influences language. **A-C** Relationship between selection for transcription factor motif integrity (x-axis) and motif association with language ability (y-axis) in (A) HAQERs, (B) HARs, and (C) random genomic regions. Each point represents one transcription factor motif. Error bars indicate ±1 standard error. Purple line (or gray for non-significant fits) shows York regression fit with 95% confidence interval (shaded); regression coefficient (*β*), chi-squared statistic (*χ*^2^), and p-values are shown. **D** Detailed view of motif effects in HAQERs colored by transcription factor family. Solid points indicate motifs with p *<* 0.05 for both positive selection and positive language association. Colored polygons show convex hulls for each transcription factor family. **E** Enrichment analysis of transcription factor families for concordant positive selection and language effects, shown as log2 odds ratios. Error bars indicate 95% confidence intervals. Solid points indicate p *<* 0.05.

Analysis of specific transcription factor families revealed striking enrichment of Homeobox and Forkhead box transcription factors associated with both enhanced binding affinity in HAQERs and improved language performance (Figure 5D). The Homeobox family displayed the strongest enrichment among all transcription factor families (odds ratio = 11.58, p-value = 2.6 × 10^−34^), followed by the Forkhead box family (which includes *FOXP2*, odds ratio = 7.28, p-value = 5.3 × 10^−6^, Figure 5E, Supplementary Table 16). These results suggest that hominin-gained binding of Homeobox and Forkhead box families within HAQERs may have played a crucial role in the evolution of human language capability.

### HAQERs regulate language-relevant brain circuits through human-specific chromatin accessibility

To determine which brain cell types are regulated by HAQERs, we analyzed their overlap with candidate cis-regulatory elements (cCREs) identified through single-nucleus chromatin accessibility profiling (snATAC-seq) of human and mouse brain cell types (*38*). We tested whether HAQERs preferentially associate with human-specific versus evolutionarily conserved chromatin accessible regions, reasoning that HAQERs should show increased overlap with human-specific regulatory elements if they provide novel functions in the human lineage.

HAQERs demonstrated significant enrichment for human-specific cCREs across brain cell types (Figure S5A), with the strongest enrichment in medium spiny neurons (MSNs, p-value = 9.8 × 10^−8^). MSNs comprise over 90% of neurons in the human striatum, a circuit that plays essential roles in vocal learning across species (*17, 39*) and was the only part of the brain robustly associated with developmental language disorders in a recent meta-analysis (*40*). HAQERs also showed enrichment around human-specific cCREs in *FOXP2*-expressing neurons (p-value = 5.3 × 10^−4^), providing independent evidence linking HAQERs to *FOXP2* regulatory networks beyond our transcription factor binding findings (Figure 5E). In contrast, HARs showed minimal enrichment for human-specific chromatin accessibility but strong enrichment for evolutionarily conserved regions (strongest in VIP neurons, p-value = 2.7 × 10^−10^, Figure S5B). These results are consistent with HAQERs providing novel regulatory functions specific to human brain development that may support language capabilities.

### Selective pressures acting on language and general cognition

Having multiple lines of evidence associating HAQERs with human language evolution, we next examined how selective pressures may have influenced language-related genetic variation over the past 20,000 years of human history using the Allen Ancient DNA Resource (AADR) (*41*). The AADR is the largest genotyped collection of ancient humans, providing harmonized genotype and metadata for each sample (like radiocarbon dating based sample ages). We identified ancient west Eurasians, then correlated their HAQER CP-PGS and the background CP-PGS with sample age (N = 3,244 individuals with remains dated between 18,775 to 150 years ago passing quality control). We see that the polygenic score for general cognition (background CP-PGS) has been subject to positive selection and has increased substantially over time (selection coefficient = 0.088, p-value = 2.1 × 10^-12^), Figure 6A. Unexpectedly, we found that HAQER CP-PGS has been stable throughout human history, indicating that ancient and modern humans carry similar numbers of language-related alleles in HAQERs (selection coefficient = -0.004, p-value = 0.71, Figure 6A).

**Figure 6:**
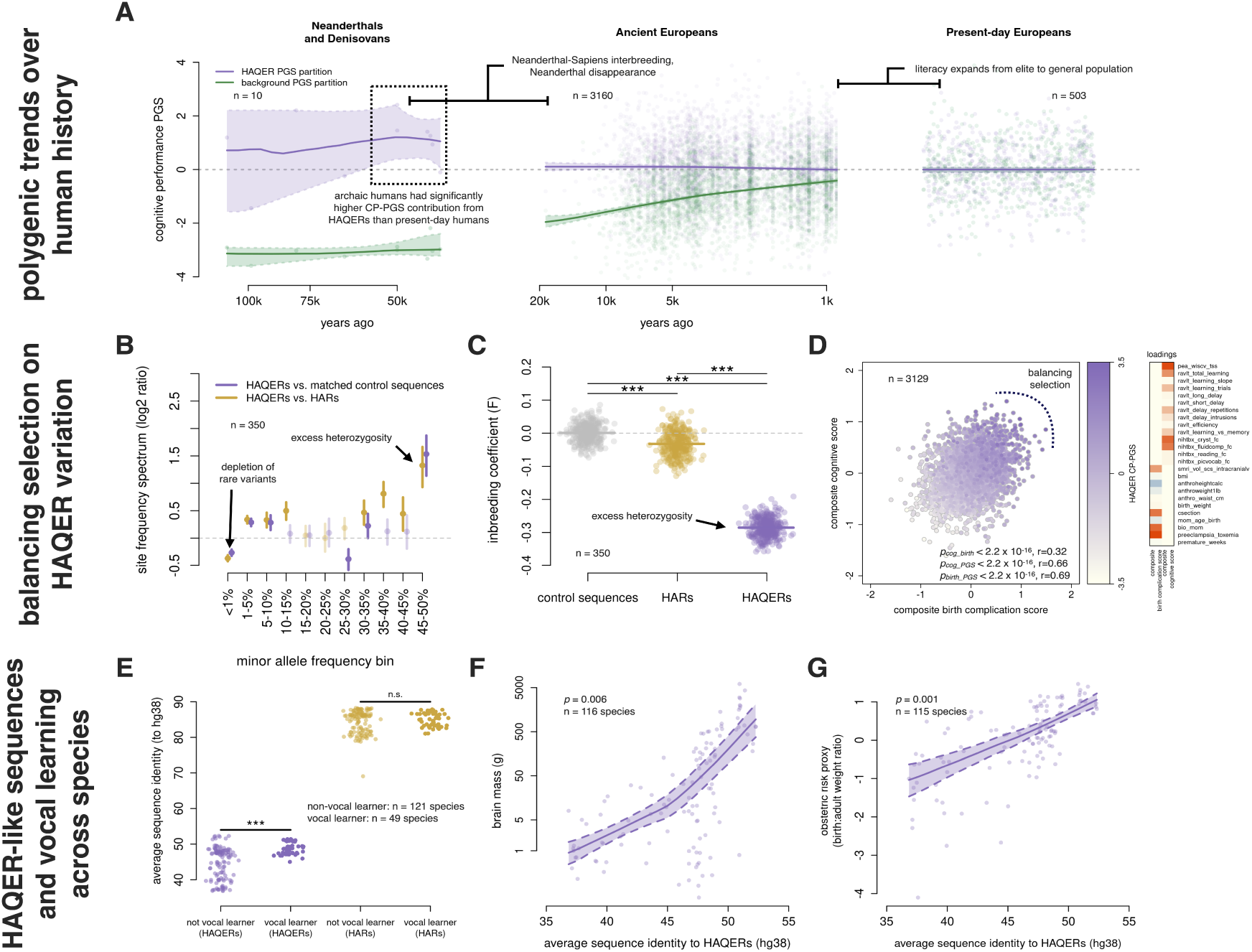
Selective pressures on human cognition and convergent evolution of vocal learning. **A** CP ES-PGS across hominin evolution. HAQER PGS (purple) and background PGS (green) plotted against sample age for Neanderthals/Denisovans, Ancient Europeans, and Present-day Europeans. LOESS fits with 95% CIs shown. **B** Site frequency spectrum comparing HAQERs to matched controls and HARs. *Log*_2_ ratio of proportion of variants across allele frequency bins; positive values indicate HAQER enrichment. Error bars: 95% bootstrap CIs. **C** Inbreeding coefficient (F-statistic) across sequence types. Lower F-statistics indicate excess heterozygosity. HAQERs show significantly lower F-statistics than HARs and control regions (*** indicates p-value *<* 0.001). **D** Composite phenotype analysis linking HAQER CP-PGS to birth complications and cognitive scores. Scatterplot: canonical correlation components colored by HAQER CP-PGS. Heatmap: variable loadings showing birth complications on component 1, cognitive variables on component 2. **E** HAQER-like and HAR-like sequence similarity scores in non-vocal learning (N = 121 species) and vocal learning (N = 49 species) mammals. Phylogenetic logistic regression statistics are shown. **F-G** HAQER-like sequence similarity correlates with brain mass (N = 116 species, **F**) and birth:adult weight ratio (N = 115 species, **G**) across mammals. LOESS fits with 95% CIs.

The presence of HAQERs in archaic humans provided a unique opportunity to describe genetically predicted cognitive traits across human species. To do this, we computed HAQER CP-PGS and background CP-PGS in archaic humans (N = 10) and compared them to ancient (N = 3,244) and modern humans (N = 503). Remarkably, the ten archaic human genomes (eight Neanderthals and two Denisovans) showed elevated HAQER CP-PGS (mean z-score = 0.91, median z-score = 1.23), while having reduced background CP-PGS (mean z-score = -3.02, median z-score = -3.01, Figure 6A). In contrast, a set of random matched control regions showed no differences in CP-PGS across groups (Supplementary Figure S4A-C, Supplementary Table 9). While this data should be interpreted with caution due to challenges applying polygenic scores across populations (*42*), the elevated HAQER CP-PGS we observed aligns with arguments that archaic humans were capable of complex language (*43–45*).

### Evidence of balancing selection from modern genomes

The stability of HAQER CP-PGS throughout human evolutionary history led us to hypothesize that HAQERs have been maintained through balancing selection. Multiple population genetic analyses in the EpiSLI sample provided support for this hypothesis. HAQERs exhibited significantly more heterozygosity compared to both HARs (t-statistic = 110, p-value = 5.3×10^-273^) and matched control sequences (t-statistic = 147, p-value = 1.6 × 10^-315^), suggesting that heterozygosity at HAQER loci provided a selective advantage (Figure 6C). Additionally, we observed an enrichment of intermediate frequency variants (MAF 30-50%) in HAQERs (Figure 6B). Both excess heterozygosity and enrichment of intermediate allele frequencies are characteristic signatures of balancing selection. Together with our ancient DNA results, these findings support a model where selective pressures prevented fixation of language-enhancing alleles in HAQERs, maintaining genetic variation at intermediate frequencies throughout human history.

### HAQERs influence prenatal brain development

The evolutionary analysis revealed a puzzling pattern: while other cognitive variants show recent positive selection, language-promoting HAQER variants have remained stable for at least the past 20,000 years of human history despite their clear cognitive benefits. This stability suggests ongoing fitness costs that counterbalance the advantages of enhanced language ability. Given HAQERs’ established role in neurodevelopment (*34*), we investigated whether these variants create pleiotropic effects on prenatal brain development that could explain their evolutionary constraints. First, we investigated temporal and cell-type enrichment for these regions to identify if they were likely to influence birth related neurodevelopmental traits. HAQERs showed broad enrichment for variants affecting prenatal brain gene expression when intersected with single-cell quantitative trait loci (scQTLs) from developing midbrain neurons (*46*). The strongest enrichment observed was at the late prenatal time point, which corresponds to when the human brain most rapidly expands (*47*) (Figure S5C). Critically, HAQERs showed no enrichment when we examined adult brain regulatory elements (*48*), confirming prenatal neurodevelopmental effects (Figure S5D). HAQERs were also significantly enriched around genomic loci associated with head circumference at birth, a proxy for brain size (*49*) (p-value = 4.4 × 10^−4^, Figure S5E).

### HAQERs link language evolution to the obstetric dilemma

The evidence for HAQER effects on prenatal brain development and head circumference at birth suggests a potential mechanism for their evolutionary stability: the obstetric dilemma. Enhanced fetal brain development may create reproductive costs through the obstetric dilemma, where increased brain size complicates birth in bipedal humans with narrow pelvises (*50,51*). To test whether HAQERs contribute to this trade-off, we analyzed brain imaging, cognitive, and birth outcome data in the ABCD cohort (*37*).

A canonical correlation analysis revealed two distinct composite phenotypic axes associated with HAQER CP-PGS, providing evidence for the obstetric dilemma that could plausibly drive the observed balancing selection (Figure 6D). The first canonical component captured variance primarily from birth complication-related variables, while the second component loaded predominantly on cognitive performance measures (including a measure of verbal language learning). Critically, both composite scores showed positive correlations with HAQER CP-PGS (*r* = 0.69 and *r* = 0.66 respectively; p-value *<* 2.2 × 10^−16^ for both), and the two composite scores were themselves positively correlated (*r*=0.32, p-value *<* 2.2 × 10^−16^). This pattern indicates that genetic variants contributing to higher HAQER CP-PGS simultaneously increase both cognition (a trait under positive selection) and birth complication risk (a trait under negative selection). The consistent positive relationship between HAQER CP-PGS and both phenotypic domains provides a plausible mechanistic explanation for the balancing selection signatures specific to HAQERs. While the correlation magnitudes should be interpreted cautiously given our optimization procedure (see Materials and Methods), the qualitative finding of antagonistic pleiotropy is robust and aligns with the evolutionary hypothesis that the cognitive benefits conferred by HAQER variants are counterbalanced by obstetric costs, maintaining genetic diversity at these loci through balancing selection. This trade-off between cognitive ability and increased birth complications is a fundamental constraint that may have shaped the evolution of human language.

### HAQER-like sequences show convergent evolution in vocal learning mammals

To test whether HAQER functions extend beyond humans, we analyzed homologous sequences across 170 non-primate mammalian species, including 49 vocal learners and 121 non-vocal learner species (*15*). Vocal learner species can acquire and modify vocalizations through experience, contrasting with species restricted to innate vocalizations. We computed genome-wide “HAQER-like” and “HAR-like” sequence similarity scores using multiple sequence alignments (*52*) and tested for associations with vocal learning ability while controlling for phylogenetic relatedness (*53, 54*).

Vocal learner species showed significantly higher HAQER-like sequence similarity compared to non-vocal learners (phylogenetic regression *β* = 1.41, p-value = 1 × 10^−4^, Figure 6E). HAQERs were also enriched around previously identified mammalian vocal learner enhancer regions (p-value = 0.028, Figure S5F) (*15*). While HARs showed a similar enrichment around vocal learner enhancer regions (p-value = 0.04), HAR sequence similarity was not associated with vocal learner classification (phylogenetic regression *β* = -0.15, p-value = 0.67, Figure 6E). Given the independent evolution of vocal learning across mammalian lineages, these results suggest HAQER-like sequences may be a fundamental genetic mechanism for complex vocal communication that has been repeatedly utilized across evolutionary history.

Remarkably, HAQER-like sequences also associated with brain size across species (phylogenetic regression *β* = 0.42, p-value = 0.006, N = 116 species) and larger relative birth weights (phylogenetic regression *β* = 0.44, p-value = 0.001, N = 115 species), mirroring the human obstetric dilemma pattern (Figure 6F-G). This convergent evidence across independent evolutionary lineages supports a link between the genetic architecture of vocal learning, brain development, and reproductive constraints. Together, these results suggests the trade-offs we observe in human language evolution may represent a broader biological phenomenon that support complex vocal communication.

## Discussion

Our evolutionary stratified polygenic score analysis identifies genomic regions that disproportionately contributed to human language evolution and continue to influence individual differences in language abilities observed in present-day humans. While previous research demonstrated that rare mutations in *FOXP2* can cause language disorders (*5, 55*), common variants in *FOXP2* show minimal association with typical language variation (*7, 8*), prompting GWAS studies and polygenic models of language-related traits (*9, 11–14, 56*). However, these models left critical questions unanswered: when did language-associated variation evolve, and how do these molecular changes influence language development? Our analysis reveals that HAQERs (*31, 34*), mostly non-coding regions that rapidly evolved before the human-Neanderthal split and represent less than 0.1% of the human genome, harbor variants with disproportionate effects on language. Individual SNPs in HAQERs carry 188 times more impact on language than variants elsewhere in the genome, while showing no association with nonverbal cognition. These potent regulatory effects distributed across thousands of small genomic elements support the polygenic architecture of human language while pointing to its specific evolutionary origins.

We find that HAQERs evolved across hominins to increase binding for Forkhead (including *FOXP2*) and Homeobox transcription factors (TFs), with motif integrity correlating with individual language scores. Supporting this regulatory mechanism, HAQERs show significant enrichment for human-specific chromatin accessible regions across brain cell types, with strong enrichment in medium spiny neurons and *FOXP2*-expressing neurons, while HARs primarily overlap with evolutionarily conserved regulatory elements. This suggests *FOXP2* influences language primarily through its regulatory networks (*57*) rather than protein-coding changes, explaining how rare mutations in a single TF can produce profound effects on language development while common variants in the same TF shows minimal associations (*7, 8*). The observed 11.58-fold and 7.28-fold enrichment of hominin-gained Homeobox and Forkhead binding sites that positively correlate with language scores in HAQERs indicates that these developmentally essential TF families (*58–60*) likely became central to human language evolution.

Cross-species triangulation provides independent support that HAQERs are functional regulatory elements for language-related traits. Consistent with our human evidence, vocal learner mammals show significantly higher HAQER-like sequence similarity than non-vocal learners after controlling for phylogenetic relationships, with parallel associations for brain size and birth weight. Importantly, HAQERs show enrichment around established mammalian vocal learning enhancer regions (*15*), consistent with previous reports of convergent evolution for complex vocal communication (*15–20*). This convergent evolution of HAQER-like sequences across vocal learning lineages provides strong independent support that these regulatory elements are fundamental genetic mechanisms for complex vocal communication.

Ancient DNA analysis reveals that while general cognitive variants show positive selection over 20,000 years, language-related HAQER variants have remained stable, suggesting balancing selection maintains language-related genetic variation. Analysis of birth outcomes, brain imaging, and cognitive data provided a potential mechanism for this unexpected evolutionary constraint: individuals with more language-related variants in HAQERs were more likely to have larger heads and birth complications, indicating trade-offs between language capability and reproductive costs. This pattern connects HAQERs to the obstetric dilemma, the evolutionary trade-off between narrower pelvises supporting upright walking and larger fetal brains enabling complex cognition (*50, 51*), potentially explaining why language-promoting variants persist at intermediate frequencies rather than reaching fixation. Further supporting this neurodevelopmental mechanism, we find that HAQERs are enriched for genetic variants associated with prenatal gene expression and head circumference at birth (*46, 49*). The prenatal brain regulatory activity of HAQERs aligns with established evidence that early developmental processes critically influence later language outcomes (*61, 62*). Intriguingly, despite substantial methodological limitations in cross-population polygenic score applications (*42*), available Neanderthal and Denisovan genomes suggest higher frequencies of language-promoting HAQER variants than modern humans, though this requires cautious interpretation. This finding needs further validation, and future research should also investigate morphological and obstetric differences between archaic and modern humans that may have enabled archaic populations to carry higher polygenic scores of language-associated alleles in HAQERs. These results point to the obstetric dilemma as an ongoing evolutionary constraint that specifically limits language-related genetic variants from reaching fixation, creating a fundamentally different selection landscape for vocal communication compared to general cognitive abilities.

These findings demonstrate how ancient regulatory innovations continue to shape human language variation through evolutionary constraints that balance cognitive benefits against reproductive costs. The independent evolution of HAQER-like sequences in vocal learning mammals reveals fundamental genetic mechanisms for complex communication that have emerged repeatedly across lineages. While general cognitive variants show positive selection over time, language variants remain stable at intermediate frequencies, suggesting that individual differences in spoken language abilities reflect ongoing evolutionary trade-offs. These evolutionary constraints reveal why individual differences in language abilities persist despite the importance of communication skills. Further investigation of HAQER regulatory networks will be essential to translate these evolutionary insights into approaches for supporting those with language disorders.

## Materials and methods summary

We developed an evolutionary stratified polygenic score (ES-PGS) approach to trace the origins of language-relevant genetic variation across 65 million years of primate evolutionary events. Language abilities were assessed through factor analysis of 17 longitudinal cognitive and language assessments from kindergarten through 4th grade in 350 children from a community-based cohort (EpiSLI) (*21*). We applied ES-PGS using cognitive performance (*28*) polygenic scores partitioned across five evolutionary annotations (*29–33*), testing whether genomic regions from specific evolutionary periods contribute disproportionately to language versus general cognitive functions.

Validation was performed across multiple independent cohorts using both common and rare genetic variants. In the SPARK autism dataset (N *>* 30,000) (*36*), we tested HAQER effects on verbal language ability and language disorder diagnoses. We analyzed rare “reversions” in SPARK whole genome sequencing data (N *>* 2,000 individuals), variants reverting from human-specific to human-chimpanzee ancestral states (*34*), reasoning that if HAQERs evolved to enhance language, reversions should impair language function. Additional validation used unrelated individuals from the ABCD cohort (N = 5,625) for spoken word recall performance, cognitive test scores, brain size, and birth traits (*37*).

To investigate molecular mechanisms supporting language evolution, we analyzed how rare genetic variants in evolutionarily significant regions affect transcription factor binding sites using position weight matrices for 633 human transcription factors from JASPAR2020 (*63*). We compared hominin-chimpanzee ancestral allele reversions versus other rare variants to detect systematic evolutionary changes in transcription factor binding associated with language phenotypes. Additionally, we looked for enrichment of evolutionarily significant regions across human-specific or conserved chromatin accessible regions (*38*), birth head circumference GWAS loci (*49*), neurodevelopmental related scQTLs datasets (*46, 48*), and mammalian vocal learning genomic regions (*15*).

Ancient DNA analysis examined selective pressures using the Allen Ancient DNA Resource (*41*), correlating HAQER polygenic scores with sample age in 3,244 ancient west Eurasians (18,775-150 years ago). We computed polygenic scores in archaic humans (8 Neanderthals, 2 Denisovans) and compared them to ancient and modern humans. Cross-species validation analyzed HAQER-like sequence similarity across 170 non-primate mammalian species (49 vocal learning, 121 non-vocal learning) (*15*) using phylogenetic regression to control for evolutionary relatedness (*53, 54*) in analyses of vocal learning, brain size, and a birth:adult weight ratio (used as a rough proxy for the obstetric dilemma) (*64*).

## General

We are grateful for the contributions of the EpiSLI cohort, their families, and the research team. We also appreciate the effort of participants, families, and research team members in ABCD, SPARK, SPARK Research Match, and Tempus. We appreciate obtaining access to genetic and phenotypic data for SPARK data on SFARI Base. We are incredibly appreciative of the open access datasets used in this study, including the researchers who put together the AADR dataset, the Zoonomia consortium datasets, and PanTHERIA. We also greatly appreciate Dr. Kristi Hendrickson and Dr. Susan Shen for valuable and encouraging discussions about this project.

## Author contributions

Conceptualization: L.G.C, T.K., J.B.T, J.J.M.; Methodology: L.G.C, T.K., T.R.T, J.B.T., J.J.M.; Investigation: L.G.C, T.K., T.R.T, JY.K, A.M., J.B.T, J.J.M.; Visualization: L.G.C, T.K., J.J.M.; Funding acquisition: J.J.M.; Project administration: D.H., S.T., M.O’B., J.C.M, J.B.T, J.J.M.; Supervision: J.C.M, J.B.T, J.J.M.; Writing – original draft: L.G.C., T.K., J.J.M; Writing – review & editing: L.G.C., T.K., T.R.T, JY.K, S.T, A.M, J.C.M., J.B.T., J.J.M

## Data and materials availability

Custom code for ES-PGS and all analyses in this paper is available here, including example code for ES-PGS that can be applied to other research problems: https://github.com/lucasgcasten/language_evolution

The EpiSLI whole genome sequencing data described here is available to qualified researches via dbGaP (study accession = phs002255.v1.p1): https://www.ncbi.nlm.nih.gov/projects/gap/cgi-bin/study.cgi?study_id=phs002255.v1.p1

SPARK genetic and phenotype data is available to qualified researchers at SFARI base:

https://base.sfari.org/

ABCD is available to qualified researchers at:

https://nda.nih.gov/abcd/request-access

1000 Genomes Phase 3 data is available at:

https://www.internationalgenome.org/data/

Allen Ancient DNA Resource:

https://reich.hms.harvard.edu/allen-ancient-dna-resource-aadr-downloadable-genotypes-present-Neanderthal

Neanderthal and Denisovan genomes:

https://www.eva.mpg.de/genetics/genome-projects/

Cross-species sequence alignment data:

https://hgdownload.soe.ucsc.edu/goldenPath/hg38/cactus447way/

PanTHERIA:

https://figshare.com/articles/dataset/Full_Archive/3531875

We used publicly available tools for data processing and analysis:

bcftools:

https://samtools.github.io/bcftools/bcftools.html

PLINK:

https://www.cog-genomics.org/plink

GCTA:

https://yanglab.westlake.edu.cn/software/gcta

LDpred2:

https://privefl.github.io/bigsnpr/articles/LDpred2.html

PRSet:

https://choishingwan.github.io/PRSice/quick_start_prset/

bedtools:

https://bedtools.readthedocs.io

Biopython:

https://biopython.org/

gonomics:

https://github.com/vertgenlab/gonomics

R:

https://www.r-project.org/

## Supplementary Materials

## Materials and Methods

### Human Subjects and IRB

All EpiSLI subjects in this study were minors who assented to participation under the University of Iowa IRB# 200511767. Secondary genetic analyses for EpiSLI subjects were approved and carried out under the University of Iowa IRB #201406727. SPARK is approved under WIRB #201703201. Approval for data collection and analysis of the SPARK Research Match study described here was conducted under the the University of Iowa IRB #201705739.

### EpiSLI Cohort

The discovery cohort of this study, referred to as EpiSLI, was originally recruited as an epidemiological study of language impairments in Kindergarteners in the state of Iowa. A more detailed description of the recruitment scheme can be found in the initial (*65*), and subsequent (*21, 66*) publications. In brief, 7,218 kindergarteners, sampled to be representative of Iowa’s population were screened for language impairment with a rapid, 40-item subset (*65*) of the TOLD-2P (*67*). Children who failed the TOLD-2P screener (i.e., have poor language ability) were over-sampled to capture a broad range of language ability in the primary cohort (n = 1,929 children). Children with autism spectrum disorder and/or intellectual disability were excluded. The entirety of the primary cohort received a more complete battery of language and cognitive assessments, while teachers and parents completed scholastic and behavioral questionnaires about the children 2nd, 4th, 8th, and 10th grades. Language and cognitive assessments included: TOLD-2P (*67*), WPPSI (*68*), a narrative story task (*69*), Woodcock Reading Mastery Tests-Revised (*70*), Word-Sound Deletion task (*71*), Random Animals-Colors task (*71*). We selected a total of 390 children from the primary cohort for whole genome sequencing. This sample represents a broad spectrum of language abilities, these 390 children were chosen by sampling from the tails of the distribution of a composite language score inward until we reached our final sample size (*8, 65*).

Behavioral assessment scores came from normed T-scores for the summary scales of the Child Behavior Checklist (CBCL) developed by ASEBA (*72*). CBCLs were completed at the 2nd grade timepoint of the study (241 of the 350 children used in this study had complete CBCL data for the analysis in Supplementary Table 2).

### Factor Analysis

An exploratory maximum-likelihood factor analysis was carried out on the language and cognitive assessment scores using the factanal function from the stats package in R (*73*), with the default “varimax” rotation. A number of factors from 1-10 were evaluated, with 7 - the largest number of factors with a nominal *χ*^2^ p-value less than 0.05 - chosen for subsequent analyses (Figure **??**A). These seven factors accounted for an estimated 62.3% of the total variance in the original data.

### EpiSLI Cognitive measures

Core language assessment scores and IQ were normalized via the general scheme described in (*65*) to account for the over-sampling of individuals with low language ability. Core language scores were corrected for age as the residual of a linear regression model using lm() in R (*73*). The age-corrected core language scores were then normalized with respect to the population using the wtd.mean() and wtd.var() functions from the Hmisc package (*74*) in R.

### Cognitive data Imputation

Most individuals for whom whole genome sequencing was generated had scores from assessments given in 2nd grade, while a minority had only 4th grade (41/390) or kindergarten (39/390) scores. Missing scores were imputed using chained random forests and predictive mean matching as implemented the missRanger R package (*75*) with 5,000 trees and pmm.k = 20.

### EpiSLI Whole Genome Sequencing

#### Sample Collection and whole genome sequencing

DNA was collected and extracted from either blood or saliva samples for 390 individuals (350 of which were used in the final analysis after quality control). DNA concentration of all sequenced samples was quantified with with Qubit 2.0 Fluorometer (Life Technologies Corporation). 390 DNA samples were sheared on a E220 Focused-ultrasonicator (Covaris) to an average size of 400 bp. Sequencing libraries were generated with a Kapa Hyper Prep kit (Kapa Biosystems) according to the manufacturer protocol. All samples had an average genome-wide coverage of at least 20X and were sequenced on a HiSeq4000 (Illumina) with 150-bp Paired End chemistry.

#### Post-sequencing QC

All sequencing data was analysed with Fastqc (v0.11.8) (*76*), where no samples failed the run modules. After genome alignment (see below), some samples were found to have significantly better coverage than others, in excess of 40x. To help control for ascertainment bias in these samples, all samples with an average coverage *>* 35x (based on initially mapped BAM) were randomly down-sampled, leading to the distribution of genome-wide coverage and average insert-size outlined in Supplementary Table 17.

#### Genome Alignment and Variant Calling

Reads were processed with bcbio (v1.1.6) (*77*) and mapped to hg19 via BWA-mem (v0.7.17) (*78*). SNVs and InDels were then called with all samples in a pool with three variant callers: GATK (v4.1.2.0) (*79*), FreeBayes (v1.1.0.46) (*80*), and Platypus (v0.8.1.2) (*81*).

Variants from each caller were filtered according to caller-specific quality metrics. For GATK, variants needed to pass all VQSR tranche thresholds. For Platypus, variants were filtered based on the goodness of fit of genotype calls, excessive region-based haplotype scores, root-mean-square mapping quality, variant quality and its ratio with read depth, low complexity sequence context, allele bias, region-based read quality, neighboring homopolymers, and strand bias. For FreeBayes, variants were filtered based on a combination of allele frequency, read depth, and overall quality. The thresholds used for filtering Platypus and FreeBayes calls were the default set by bcbio. An ensemble of all three callers was generated and used in all subsequent analyses in order to achieve improved specificity in the detection of rare variants. All variants in the ensemble callset were called by GATK as well as either Platypus OR FreeBayes. The majority of variants ( 87%; 25,775,508) were called by all three callers, while a minority were called by GATK and only one other caller ( 13%; 3,937,146). All variants were filtered to have a QUAL and individual quality score ≥ 20. Qualified researchers can access the individual level genotypes from dbGaP at (study accession = phs002255.v1.p1): https://www.ncbi.nlm.nih.gov/projects/gap/cgi-bin/study.cgi?study_id=phs002255.v1.p1

#### Final sample QC

Starting from the initial sample size of 390, a total of 14 samples were flagged for removal from all association analyses because of population stratification. Specifically, these samples did not cluster with 1,000 Genomes Europeans (*82*), or were more than 3 standard deviations away from the rest of the EpiSLI cohort based on the top 10 multidimensional scaling components calculated from SNPs found at or above a 0.05 minor allele frequency. An additional 26 samples were dropped due to relatedness or limited phenotypic data, leaving a final sample size of 350 unrelated European individuals with complete data for all genomic analyses.

#### Variant Annotations

Variants were annotated with the Ensembl Variant Effect Predictor tool (VEP v109) (*83*). Reference population allele frequencies came from 1000 Genomes Phase 3 samples (*82*) and GnomAD (*84*), rsIDs of variants were dbSNP (*85*) (v151), these additional annotations were added using VCFanno (*86*) (v20190119).

### Polygenic Scores (PGS)

#### Genotypes for PGS

To derive a set of SNPs suitable for PGS analysis, we merged our dataset with 1000 Genomes Europeans to use as a reference sample. Based on widely used recommendations (*87*), we extracted SNPs with a minor allele frequency ≥ 1%, Hardy-Weinberg equilibrium p-value *>* 1 × 10^-6^, and a missingness rate *<* 2% in both samples (leaving 7,719,665 SNPs total for PGS calculation).

#### PGS Calculation

LDpred2 was used to calculate a genome-wide PGS for all traits with the infinitesimal model using the provided UK Biobank LD reference panel and HapMap3+ variant set (*88*). PGS were calculated using GWAS summary statistics for neurodevelopmental and psychiatric traits: ADHD (*89*), addiction (*90*), alcohol dependency (*91*), Alzheimer’s (*92*), Autism (*93*), Anorexia (*94*), bipolar disorder (*95*), cannabis use disorder (*96*), depression (*97*), epilepsy (*98*), insomnia (*99*), neurodevelopmental conditions (*100*), PTSD (*101*), schizophrenia (*102*), and Tourette’s (*103*). GWAS summary statistics for cognitive traits included: cognitive performance (*104*), educational attainment (*105*), executive functioning (*106*), and the “g Factor” (*107*). Additional PGS were calculated for the following behavioral and socioeconomic status related traits: childhood aggression (*108*), antisocial behavior (*109*), empathy (*110*), the BIG5 personality traits (*111*), income (*112*), and the Townsend Deprivation Index (a measure of material deprivation) (*113*). PGS were also calculated for brain structural (*114, 115*) and functional connectivity phenotypes (*116*). Finally, we computed PGS for miscellaneous traits: left-handedness (*113*), height (*117*), childhood trauma (*118*), and vocal pitch (*119*). It is important to note we did not compute PGS in EpiSLI for reading based traits, because our sample was part of the discovery cohort of the largest reading related GWAS to date (*9*). Associations for all of these PGS with our language factors can be found in Supplementary Table 3.

To account for population stratification, we corrected PGS for the first 5 genetic principal components. PGS were normed to the 1000 Genomes Europeans reference sample.

#### ES-PGS Calculation

We developed evolutionary stratified polygenic scores (ES-PGS) to identify genomic regions from specific evolutionary periods that disproportionately contribute to modern phenotypic variation. ES-PGS partitions polygenic score effects based on evolutionary annotations, testing whether variants from particular evolutionary epochs show disproportionately strong associations with traits compared to the broader genome.

The method uses three components: (1) annotation-specific PGS calculated from variants within evolutionary regions of interest, (2) background PGS calculated from all remaining genome-wide variants, and (3) matched control PGS from biologically similar regions to test specificity. Matched control regions were generated by sampling 1,000 random genomic segments for each annotation region (e.g., 10 annotation region would produce 10,000 random regions for matching), these regions are then matched for chromosome, size, GC content, repeat content, distance to nearest gene, and overlap with promoter or coding sequences. Control regions were required to be *geq* 100kb from any annotation region to ensure independence.

We implemented ES-PGS using nested model comparisons. The reduced model tests background genomic effects and matched control effects:

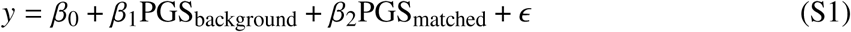

The full model adds the annotation-specific term:

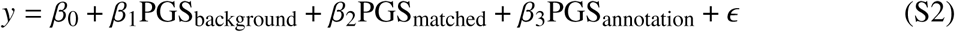

Where *y* represents the phenotype (e.g., language factor scores), PGS_background_ includes all variants except those in the focal annotation, PGS_matched_ represents the matched control regions, and PGS_annotation_ contains only variants within the evolutionary annotation. An ANOVA comparing these models tests whether the annotation contributes significantly beyond background genetic effects and matched controls. Additional covariates (genetic principal components, demographics) can be incorporated as needed.

PRSet (*27*) was used to calculate stratified PGS using human genome annotations in BED formatted files, computing a clumping and thresholding based ES-PGS for each annotation and a background PGS for the rest of the genome. Background regions for each annotation were identified using the “complement” function from bedtools (v2.26.0) (*120*). Matched control regions were matched using the nullranges R package (*121*). As recommended by the authors of PRSet, we used a p-value threshold of 1 and the 1000 Genomes Europeans as the LD reference (*82*). We corrected for population stratification using the same steps as described in the genome-wide PGS analysis.

We focused our analysis on genome annotations related to primate and human evolution, annotations included: primate ultra conserved regions (primate UCEs, Supplementary Data 5 from the reference) (*29*), great ape lineage accelerated regions (*30*)(Supplementary Data S1 filtered to “Hominidae” from the reference), Human Ancestor Quickly Evolved Regions (HAQERs, Supplementary Table 68 from the reference) (*31*), Human Accelerated Regions (HARs) (*32*), and ancient human selective sweep regions (Supplemental File S2 from the reference) (*33*). All annotations were converted to BED format and lifted over to match genome builds when necessary, using the UCSC liftOver tool (*122*). All annotation coordinates and matched regions can be found in our repository: LINK XX

#### ES-PGS replication in SPARK

To validate our ES-PGS results in a separate sample we used the SPARK cohort, a large genetic study of individuals with autism and their family members (*36*). For the replication, we utilized imputed SNP array data that we have previously described (*123*) to compute an ES-PGS using the same workflow as in our EpiSLI sample. The self-reported language and psychiatric diagnosis phenotypes, came from a questionnaire as part of a previously described research match where we had *>* 1,000 adults with autism or parents of children with autism complete an online language battery (*124*). We grouped all self-reported language-related diagnoses (language impairment, dyslexia, hearing impairment, stuttering, and speech impediments), and psychiatric diagnoses together (major depression, generalized anxiety disorder, bipolar, schizophrenia, substance abuse, OCD, and Tourette’s). Given the large number of people without any language or psychiatric diagnosis, we used zero inflated Poisson regression models to determine the relationship of the HAQER CP-PGS with having any language or psychiatric diagnosis, as well as the number of language or psychiatric diagnoses. We used the log-ratio test to determine if adding the HAQER CP-PGS term significantly improved prediction accuracy in the zero inflated Poisson regression models. All other phenotypes came from the SPARK December 2014 phenotype release. All analyses included age, sex, background CP-PGS, HAQER matched control CP-PGS, and the first 5 genetic principal components as covariates.

#### SPARK rare ancestral reversion analysis

To explore the effects of rare genetic variation in evolutionary significant regions on language ability, we used the whole genome sequencing data from the previously described SPARK cohort (max N = 11,545) (*36*). Briefly, we merged the whole genome sequencing data provided by SPARK (WGS batches 1-4) using bcftools, then used the same processing pipeline as we did for the EpiSLI cohort: filtered to variants with a QUAL and individual quality score ≥ 20, and annotated the variants with VEP (v109) (*83*). To identify rare variants we then filtered variants in both datasets to have a maximal reference population allele frequency *<* 1% and an allele frequency *<* 1% in SPARK. Ancestral alleles were identified using those provided in the original HAQER manuscript (*34*). We identified all reversion variants within 10Kb of HAQERs from the updated set (*31*), HARs (*32*), or random non-coding (RAND) (*34*) sequence and counted the number of rare ancestral reversions each sample had in each of these elements. We removed outlier samples who had *>* 2.5 median absolute deviations away from the median value for either HAQER, HAR, or random reversions, but had consistent phenotypic associations even when including these outliers in the analysis. We then used these reversion counts for association with speech and language phenotypes in SPARK.

### Transcription Factor Analysis

#### Variant selection and annotation

We analyzed transcription factor binding sites in three distinct genomic contexts: Human Accelerated Regions (HARs), Human-Accelerated Quickly Evolved Regions (HAQERs), and matched random genomic regions (RAND, taken from (*31, 34*)). Position weight matrices (PWMs) for 633 human transcription factors were obtained from the JASPAR2020 database (*63*), with pseudocounts adjusted according to base frequency distributions.

Variants were filtered using strict quality control criteria as described above. For this analysis, we retained only biallelic single nucleotide variants (SNVs) located within feature boundaries that exhibited minor allele frequencies below 1% across all reference populations provided by VEP (*83*) and a minor allele frequency below 5% in our smaller EpiSLI sample. Complete great ape allele information was required for each variant, including data from the human reference (hg19), Neanderthals and Denisovans, Chimpanzee, Bonobo, Gorilla, and Orangutan genomes (*34, 125–128*). Briefly, archaic hominin genotype data came from VCFs with genotypes for the three high coverage Neanderthals (Altai (*125*), Chagyrskaya (*126*), and Vindija (*127*)) and one Denisovan (*128*) produced from high-coverage while genome sequencing were downloaded from the Max Planck Institute for Evolutionary Anthropology website and merged with our data. The final dataset comprised genotype information from 15,729 rare variant sites across 350 individuals, with corresponding variant annotations.

#### Reversion Status Determination

To characterize the evolutionary trajectory of variants, we developed a machine learning approach to impute hominin (human reference and most common allele from Neanderthal and Denisovan genomes matched) to human-chimp ancestral reversion status where direct determination was not available. We implemented an elastic net regression model (*α* = 0.9) using the glmnet package in R (*129*), incorporating great ape allele states as predictors of reversion status as given in (*34*). To address class imbalance, we applied weights to ensure equal representation across sequence context types (HAQER, HAR, random). Reversion status was assigned using a probability threshold of 0.95 (i.e., at least a 95% predicted probability of being a human-chimp ancestral reversion), with known states preserved for training data.

#### Sequence and Motif Analysis

For each variant, we extracted 51-base pair genomic windows centered on the variant position from the human reference genome (hg19). Alternative sequences were generated by substituting variant alleles into the reference background. We then calculated maximal motif scores for both reference and alternative sequences across all JASPAR2020 human transcription factor motifs. Scores were computed on both forward and reverse complement strands, with the maximum score retained for reference and alternate alleles of each variant-motif pair.

#### Language Association Analysis

Core language ability was assessed using the F1 measure as described above in the factor analysis. We computed burden scores for each transcription factor motif by combining variant effects weighted by genotype status of reversion sites. Linear regression models were used to estimate the associations between individual context-specific (aggregate) motif scores and language ability.

#### Transcription Factor Motif Score Selection Analysis

For each transcription factor, the (Z-scaled) difference in reference and alternate allele motif scores was modeled as a linear function of variant sequence context (HAQER, HAR, or random) and a binary reversion status indicator. Separate reversion effects were estimated for HAR, HAQER, and random region variants as a sequence context by reversion interaction term. Estimated beta coefficients from these terms, as well as their standard errors, were extracted for use in downstream analyses.

#### Joint Selection-Language Enrichment Analysis

We used York regression analysis (*130*) to examine relationships between hominin-divergent motif integrity and language-related effects. This approach accounts for uncertainty in both variables (i.e., language association betas and hominin-divergent binding effect betas). Prior to the analysis, the sign of the hominin-divergent binding binding betas were flipped such that positive values indicate human-gained motif integrity scores (i.e., when reversions to the human-chimp ancestral allele tend to decrease motif scores). York regression betas, their standard errors, a Chi-squared goodness of fit statistic and its p-value were extracted to interpret the significance of the overall relationship between TF motif integrity (i.e., motif score) and motif score effect on individual differences in language ability, for each sequence context. Individual TF motifs of interest are those with nominal significance (p *<* 0.05) for both selection for motif integrity and for positive association of aggregate motif integrity with higher F1 core language scores. Transcription factors were classified into families using InterPro annotations (*131*) of representative families found to be significantly associated with a reversion effect in either direction (using www.string-db.org) (*132*). To identify transcription factor families showing convergent patterns of hominin selection and language association, we performed 2×2 Fisher’s exact tests comparing the proportion of motifs with concordant effects (hominin gained binding AND positive language association vs. all other combinations) across families. Odds ratios with 95% confidence intervals and p-values were computed for each TF family.

#### Ancient DNA

DNA data and sample age information for ancient *Homo sapiens* came from the Allen Ancient DNA Resource (AADR) version 54. We downloaded the publicly available EIGENSTRAT formatted files and converted them to PLINK format using the EIGENSOFT tool. We then merged the ancient genomes with our EpiSLI and 1000 Genomes Europeans dataset to ensure we were using comparable SNPs for our ES-PGS selection analysis as we did in our discovery sample. We identified ancient west Eurasians using the same criteria as a recent large-scale selection analysis (*133*). Briefly, we filtered to samples found between longitude 25W and 60E and latitude 35N to 80N, samples passing quality control with an assessment labeled as “PASS”, and sample ages *>* 0 but *<* 20,000 years old.

We then computed ES-PGS for CP in HAQERs using the same methodology as we did in the EpiSLI, SPARK, and ABCD samples for use in our polygenic selection analysis. Given the challenges of accounting for population structure in ancient DNA, we opted to use a LMM based approach instead of a traditional PC based approach as recommended by Akbari et al., 2024 (*133*). With the ancient west Eurasian subsample, we then identified independent SNPs using 1000 Genomes as the LD reference with PLINK’s “–indep-pairwise” function (window size = 1000bp, step size = 1bp, r^2^ threshold = 0.05, MAF ≥ 5%) (*134*). Next, we identified samples and SNPs with low missingness for GRM calculation (samples missing *<* 50% of independent SNPs, and SNPs missing in *<* 10% of those samples). Finally, we computed the genetic relatedness matrix (GRM) with GCTA (*135*) with the QC passing samples and SNPs and removed duplicate/twin samples for subsequent analysis (GCTA “grm-cutoff” of 0.9).

We then used the 3,244 QC passing samples and the GRM based on the 12,146 QC passing SNPs for the ES-PGS analysis. We implemented the LMM based polygenic selection analysis with the gaston (*136*) R package (lmm.aireml function), allowing us to account for the GRM which reflects population structure and relatedness of the sample. We used log10(sample age) as the outcome variable and HAQER CP-PGS and background CP-PGS as the independent variables (similar to the ES-PGS analysis we used in our other samples).

Additionally, we computed HAQER CP-PGS in the 10 available that are part of this release of the AADR. We compared these archaic hominins to ancient humans (described above), and 1000 Genomes Europeans, all data was processed together using the same set of SNPs to ensure polygenic scores would be comparable.

#### Detecting balancing selection in modern genomes

To detect signatures of balancing selection, we analyzed the WGS data we generated in EpiSLI. First, we subset to regions of interest (HAQERs, HARs, RAND, and HAQER matched control sequences). Then we computed a site frequency spectrum (SFS) for each sequence class, calculating the proportion of variants in each minor allele frequency bin. We compared HAQERs SFS to the SFS of HARs, RAND, and matched control sequences, to identify whether HAQERs had a relative enrichment of intermediate frequency variants -which can indicate balancing selection (or ongoing selection). To create 95% confidence intervals for HAQERs we generated 1,000 random bootstrap samples of HAQERs (sampled with replacement) and computed the allele frequency proportions within in each sampled HAQER set, These confidence intervals allowed us to more confidently determine whether there was enrichment or depletion across MAF bins in HAQERs when compared to other sequence types.

Next, to more formally test our balancing selection hypothesis we identified common independent SNPs in these regions using PLINK (*134*), using the “–indep-pairwise” function (window size = 200bp, step size = 50bp, r^2^ threshold = 0.5, MAF ≥ 5%) (*134*). This identified common independent SNPs in HAQERs, HARs, RAND sequences. We then computed individual level method of moments F-coefficients for each class of variation using the “–het” function in PLINK (*134*), which derives statistics based on the expected number of homozygotes and observed homozygotes (with more negative values indicating there are more heterozygotes than expected). We compared values between classes using paired t-tests to determine if there was excess heterozygosity in HAQERs, a signature of balancing selection.

#### ES-PGS analysis in ABCD

To explore the effects of HAQER CP-PGS in prenatal development we analyzed the ABCD cohort, a large longitudinal study of adolescent development with genetic, cognitive, brain imaging, and developmental phenotypes (*137*). All phenotypes used in ABCD came from the v4.0 data release. Similar to the SPARK replication analysis, we utilized imputed SNP array data that we have previously described (*123*) to compute ES-PGS using the same workflow as in our EpiSLI and SPARK samples. We computed genetic relatedness using GCTA (*135*) in the merged ABCD and SPARK dataset. Using the relatedness matrix, we identified unrelated individuals for ES-PGS analysis with brain imaging, cognitive, and birth phenotypes (genetic relatedness *<* 0.05). All evolutionary stratified polygenic scores were adjusted for genetic principal components, similar to the SPARK and EpiSLI cohorts. We included age, sex, and genetic principal components as covariates in the ES-PGS analyses.

To investigate potential evolutionary trade-offs underlying the balancing selection observed in HAQERs, we examined the relationship between HAQER CP-PGS and both cognitive performance and birth complications in ABCD. We employed a two-stage approach to address confounding and maximize signal detection. First, we applied a modified Remove Unwanted Variation using residuals (RUVr) approach (*138*) to remove variance components in the phenotypic data that were orthogonal to the HAQER CP-PGS signal. Specifically, we identified and removed k = 15 principal components from the phenotype matrix that captured variance attributable to measured confounders (age, sex, ancestral background from genetic principal components) and unmeasured technical or biological factors not associated with HAQER CP-PGS. This denoising step preserved only phenotypic variance potentially related to our polygenic score of interest while removing confounding sources of variation. Second, we performed canonical correlation analysis (CCA) (*139*) on the RUVr-adjusted phenotypes to identify composite phenotypic dimensions that maximally correlated with HAQER CP-PGS. The CCA simultaneously analyzed two phenotypic domains (each having a block): cognitive performance measures (including memory, executive function, and processing speed tasks) and birth complication-related variables (including cesarean section delivery, intracranial volume measured from MRIs, and other obstetric risk factors). Sparsity constraints (*λ* = 0.75) were applied to the phenotypic loadings to improve interpretability. While we present correlation statistics for completeness, we acknowledge these may be inflated due to the optimization procedure; our primary inference focuses on the qualitative pattern of associations rather than specific effect sizes.

#### HAQER enrichment analysis

To determine enrichment between loci of interest (prenatal scQTLs, birth head circumference GWAS loci, and mammalian vocal learning enhancers) with our evolutionary regions of interest (HAQERs, HARs, and random sequence), we used the “intervalOverlap” function from gonomics (*140*). This allowed us to compute expected versus actual overlaps (based on size of the human genome and the provided annotations) providing an enrichment p-value for each region.

#### Evolutionary annotations

Given the two sets of HAQER annotations available (*31, 34*), created using different reference genomes and sequencing methodology we opted to use a conservative set of HAQERs for enrichment analyses to ensure any associations were not due to artifacts. The conservative set of HAQERs was identified by overlapping regions in the two separate HAQER sets using the bedtools “intersect” function (*120*), leaving nearly 900 overlapping HAQER regions. We included the set of HARs and random sequence (matched to basic biological properties in HAQERs like size and non-coding sequence) from the original HAQER paper (*34*) for all enrichment analyses, allowing us to directly compare annotations. Similar to the other genomic analyses, we restricted analysis of all annotations to autosomal regions.

#### Defining human-specific and cell-type specific chromatin accessible regions

To identify whether HAQERs provided humans with novel regulatory mechanisms in specific neural cell-types, we examined data from snATAC-seq of human and mouse brains (*38*). To identify human-specific and cell-type specific cCREs, we used Suppelemental Table 6 provided by the authors to get hg38 coordinates of cCREs and filtered to “humanSpec” cCREs using data provided in Supplementary Table 20. Similarly, to identify human-mouse conserved cCREs, we used Suppelemental Table 6 provided by the authors and filtered to “CA cons” cCREs using data provided in Supplementary Table 20. We added a 100Kb flank around all chromatin accessible regions and merged overlapping regions using the bedtools “merge” function, allowing us to test whether evolutionary annotations are enriched around cell-type specific chromatin accessibility regions.

#### Defining neurodevelopmental scQTLs

To determine if HAQERs, HARs, and RAND sequences significantly influence gene expression in prenatal and postnatal brains, we leveraged scQTLs from two studies. (1) a study of stem cell derived neurons from *>* 200 individuals, meant to mimic early prenatal cells in the midbrain (*46*) and (2) a study of nearly 400 postmortem adult brains from psychENCODE2 (*48*). For scQTL study (1), we defined scQTLs as all common variants (MAF *>* 5%) with a p-value *<* 5 × 10^-4^, for scQTL study (2) we utilized all provided scQTLs defined as significant by the authors downloaded from https://psychscreen.wenglab.org/psychscreen/downloads.

#### Defining birth head circumference regions

We identified birth head circumference associated genomic loci with a genome-wide suggestive p-value (p-value *<* 5 × 10^-5^) and a MAF *>* 5%, using the largest GWAS available to date (*49*). Given the influence of LD on human traits, we added a 100Kb flank around all associated SNPs and merged overlapping regions using the bedtools “merge” function, allowing us to test whether evolutionary annotations are enriched around associated SNPs.

#### Defining mammalian vocal learning enhancer regions

We gathered previously established mammalian vocal learning enhancer regions from a recent cross-species analysis (*15*), which identified 50 open chromatin regions in the motor cortex strongly associated with vocal learning. Since these regions were provided in mouse genome coordinates (mm10), we lifted these over to hg19 and we added 1Mb flanks around these enhancer regions which were then used to test whether evolutionary annotations are enriched near established vocal learning enhancers.

#### Convergent evolution of vocal learning analysis

To test for evidence of convergent evolution of “HAQER-like” sequences in vocal learning species, we utilized a large dataset of *>* 400 species with whole genome data aligned to the human reference genome (hg38) (*52*). We parsed the alignments with Biopython (*141*) to subset to regions of interest. For each species and sequence type (HAQERs or HARs), we computed a sequence similarity (using the number of bases matching the human reference genome in these regions). We then used this “HAQER-like” sequence similarity to predict vocal learning status in 170 non-primate species that have been previously described (*15*). To determine statistical significance and account for species relatedness we used phylogenetic logistic regression (*53*) with the phylolm package (*54*) in R. We did similar analyses for brain size and birth:adult weight ratio using species level phenotype data from the PanTHERIA dataset (*64*).

#### Statistical analysis

All statistical analysis was done in R (version 4.3.1) (*73*).

#### Approach to multiple testing correction and triangulating evidence

To address multiple testing concerns, we employed False Discovery Rate (FDR) correction for analyses testing more than 20 hypotheses, allowing us to limit type 1 errors (false positives). This was applied to the EpiSLI CBCL mental health score analysis, genome-wide PGS analysis, and the ES-PGS analysis where we conducted many statistical tests.

We adopted a comprehensive approach to validate our key findings through multiple lines of evidence. We established confidence in our results through: (1) replication across independent cohorts (e.g., EpiSLI, SPARK, and ABCD); (2) identifying convergent evidence across different analytical approaches (e.g., common variant, rare variant, and transcription factor binding analyses); (3) demonstration of specificity, showing HAQERs’ associations with language but not nonverbal IQ; (4) evolutionary support from ancient DNA and cross-species analyses; and (5) mechanistic support through analysis of transcription factor binding, enrichment for human-specific and cell-type specific chromatin accessible regions, and enrichment of variants influencing prenatal gene regulation. This multi-faceted approach allowed us to distinguish robust biological signals from statistical noise, even when some individual analyses showed moderate statistical significance. Our findings’ consistency across diverse data types, species, and analytical methods provides stronger evidence than would be achieved through any single statistical test, regardless of its p-value.

## Supplementary Figures

**Figure S1:**
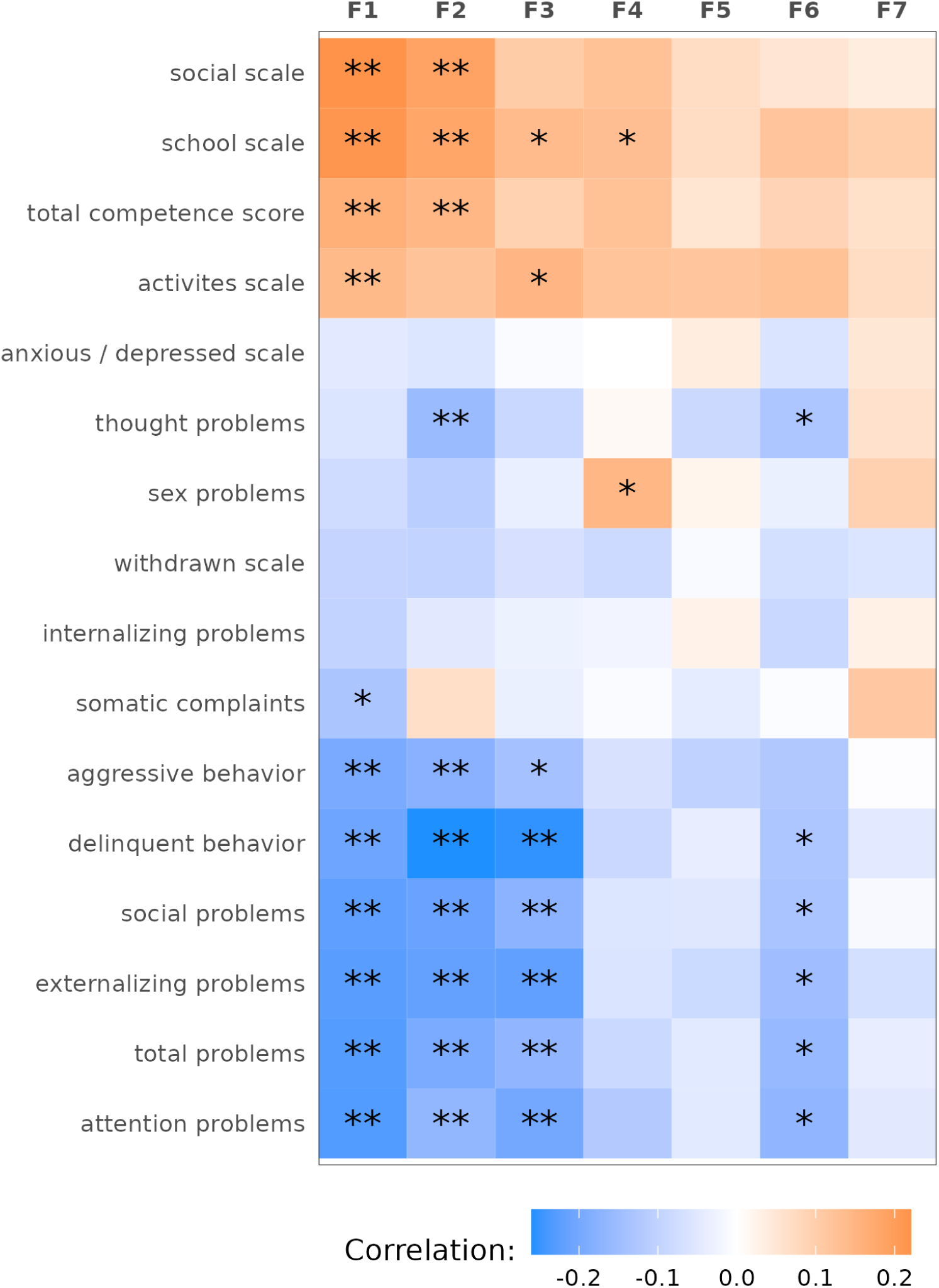
Mental health correlates of EpiSLI factors. Heatmap of Pearson correlation coefficients (*r*) of CBCL scales with EpiSLI factor scores (N = 241 individuals). Color indicates the association between that evolutionary annotation and the EpiSLI factor score (orange = positive correlation, blue = negative correlation. “**” indicates FDR *<* 0.05, and “*” indicates unadjusted p-value *<* 0.05.

**Figure S2:**
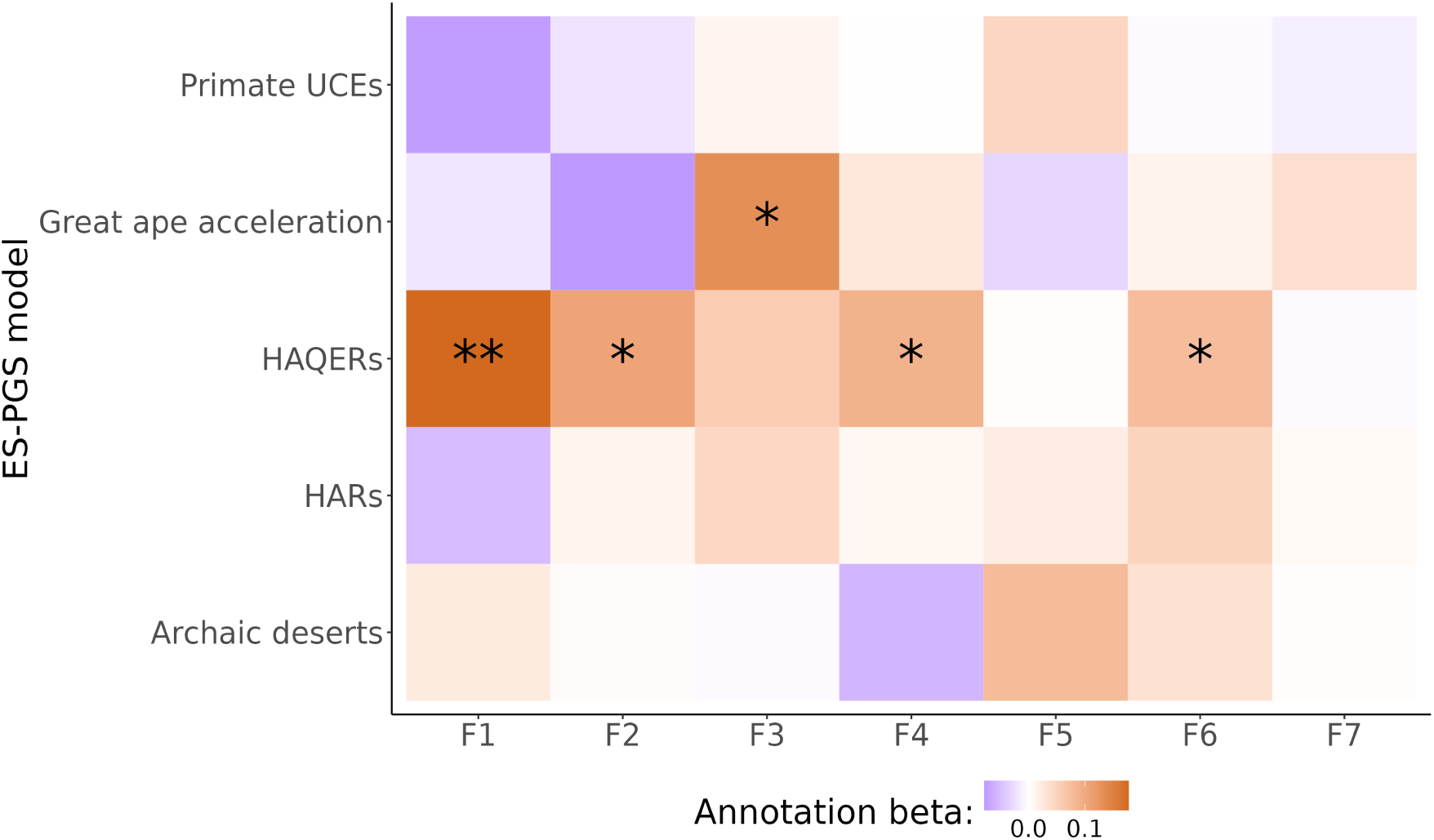
Evolutionary history of EpiSLI factors. Heatmap of ES-PGS *β*’s stratifying the CP-PGS based on evolutionary annotations. Color indicates the association between that evolutionary annotation and the EpiSLI factor score (orange = positive ES-PGS *β*, purple = negative ES-PGS *β*). “**” indicates FDR *<* 0.01, and “*” indicates unadjusted p-value *<* 0.05.

**Figure S3:**
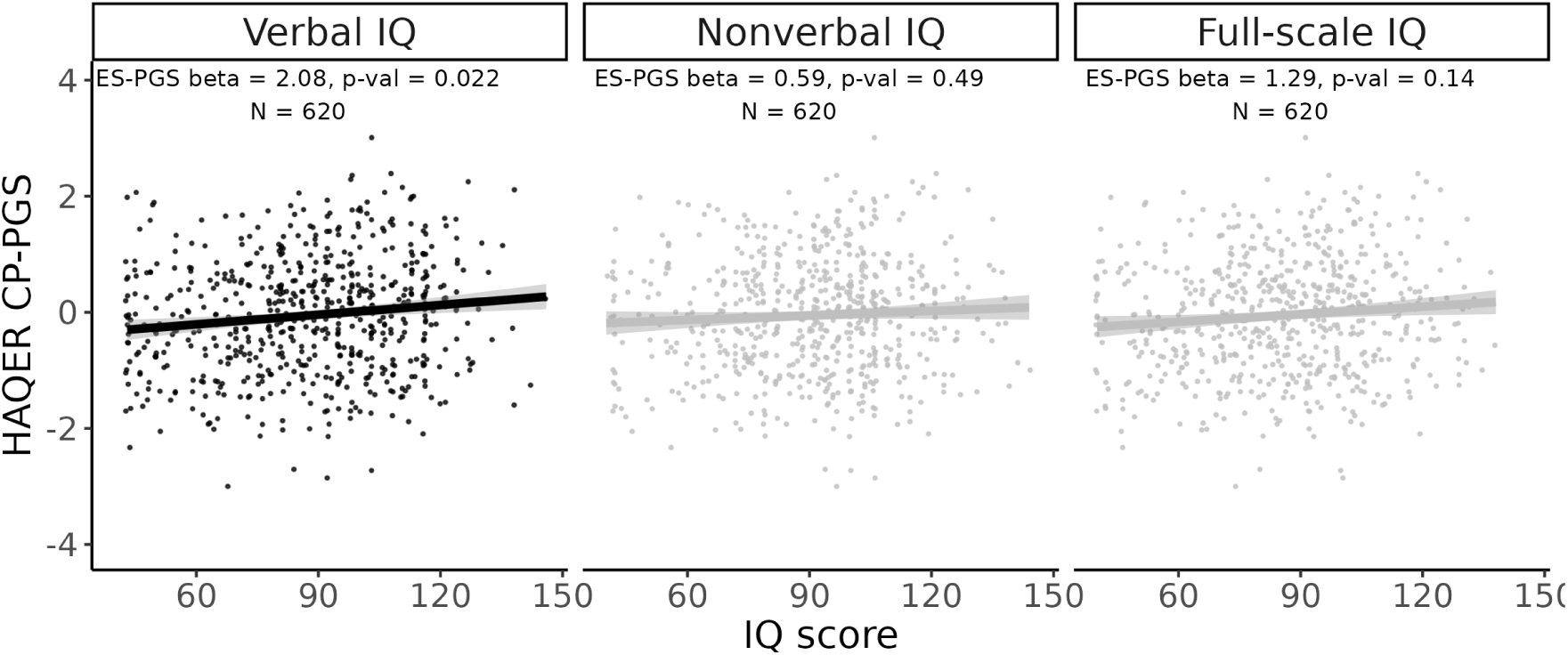
HAQER CP-PGS is associated with verbal IQ in an independent cohort. ES-PGS results of HAQER CP-PGS and clinical IQ assessment scores from the SPARK dataset (N = 620).

**Figure S4:**
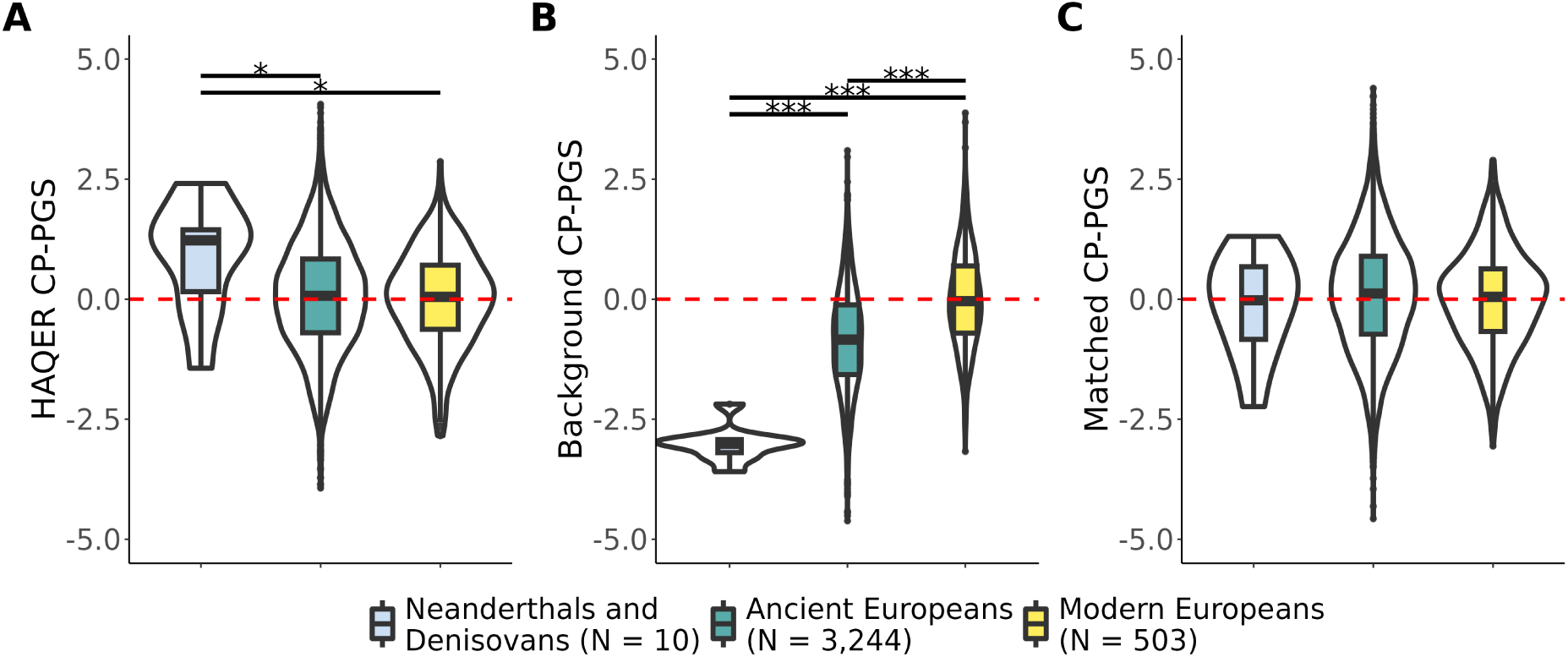
ES-PGS differences across human history. **A-C** Distribution of HAQER CP-PGS (**A**), background CP-PGS (**B**), or random matched control regions (**C**) across archaic humans (neanderthals and denisovans from the AADR), ancient anatomically modern humans (AADR), and modern Europeans (1000 Genomes dataset). “*” indicates a p-value of *<* 0.05 from a t-test comparison, and “***” indicates p-value *<* 0.001.

**Figure S5:**
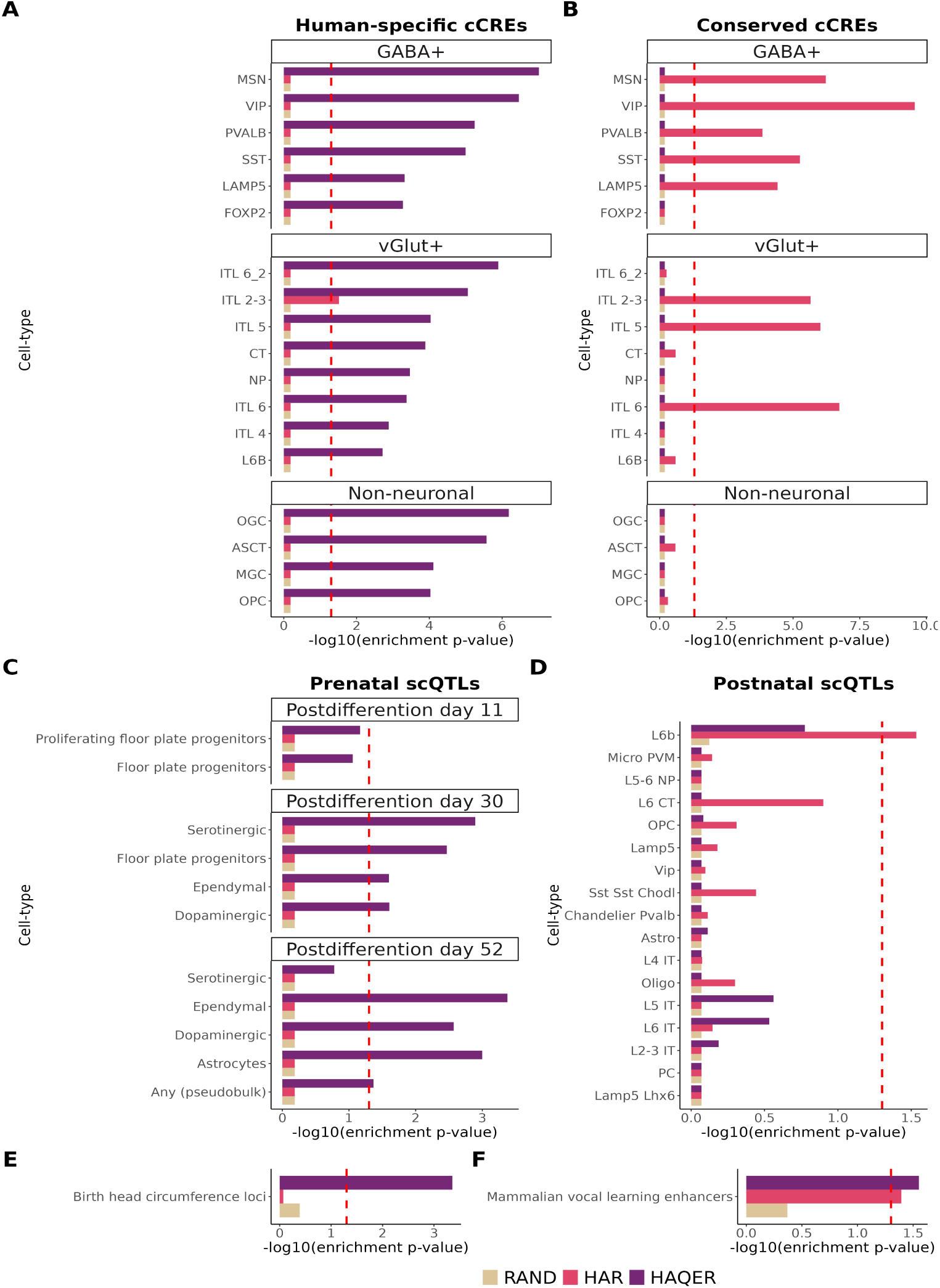
HAQERs influence prenatal brain development. **A-B** Enrichment of HAQERs, HARs, and RAND for brain cCREs from human-specific (**A**) or human-mouse conserved (**B**) elements. **C-D** Enrichment for regulatory variants (scQTLs) across timepoints and cell-types in newly differentiated neurons (**C**) or adult post mortem brains (**D**). **E** Enrichment for common SNPs (MAF *>* 5%) associated with head circumference at birth. **F** Enrichment for enhancer regions associated with vocal learning across mammals (Wirthlin et al., 2024).

